# GRA47 and GRA72 are *Toxoplasma gondii* pore-forming proteins that influence small molecule permeability of the parasitophorous vacuole

**DOI:** 10.1101/2023.11.15.567216

**Authors:** Mebratu A. Bitew, Pablo S. Gaete, Christopher Swale, Parag Maru, Jorge E. Contreras, Jeroen P. J. Saeij

## Abstract

*Toxoplasma gondii*, a medically important intracellular parasite, uses GRA proteins, secreted from dense granule organelles, to mediate nutrient flux across the parasitophorous vacuole membrane (PVM). GRA17 and GRA23 are known pore-forming proteins on the PVM involved in this process, but the roles of additional proteins have remained largely uncharacterized. We recently identified *GRA72* as synthetically lethal with *GRA17*. Deleting *GRA72* produced similar phenotypes to *Δgra17* parasites, and computational predictions suggested it forms a pore. To understand how GRA72 functions we performed immunoprecipitation experiments and identified GRA47 as an interactor of GRA72. Deletion of *GRA47* resulted in an aberrant ‘bubble vacuole’ morphology with reduced small molecule permeability, mirroring the phenotype observed in *GRA17* and *GRA72* knockouts. Structural predictions indicated that GRA47 and GRA72 form heptameric and hexameric pores, respectively, with conserved histidine residues lining the pore. Mutational analysis highlighted the critical role of these histidines for protein functionality. Validation through electrophysiology confirmed alterations in membrane conductance, corroborating their pore-forming capabilities. Furthermore, Δ*gra47* parasites and parasites expressing GRA47 with a histidine mutation had reduced *in vitro* proliferation and attenuated virulence in mice. Our findings show the important roles of GRA47 and GRA72 in regulating PVM permeability, thereby expanding the repertoire of potential therapeutic targets against *Toxoplasma* infections.

**IMPORTANCE:** *Toxoplasma gondii* is a parasite that poses significant health risks to those with impaired immunity. It replicates inside host cells shielded by the parasitophorous vacuole membrane (PVM), which controls nutrient and waste exchange with the host. GRA72, previously identified as essential in the absence of the GRA17 nutrient channel, is implicated in forming an alternative nutrient channel. Here we found that GRA47 associates with GRA72 and is also important for the PVM’s permeability to small molecules. Removal of GRA47 leads to distorted vacuoles and impairs small molecule transport across the PVM, resembling the effects of GRA17 and GRA72 deletions. Structural models suggest GRA47 and GRA72 form distinct pore structures, with a pore-lining histidine critical to their function. *Toxoplasma* strains lacking GRA47, or those with a histidine mutation, have impaired growth and reduced virulence in mice, highlighting these proteins as potential targets for new treatments against Toxoplasmosis.

## Introduction

*Toxoplasma gondii* is an obligate intracellular parasite that infects a wide range of warm-blooded hosts, including humans. It poses severe health risks to immunocompromised individuals and can lead to congenital complications in pregnant women (1). To establish a successful infection, *Toxoplasma* has evolved intricate strategies to invade and manipulate host cells (2). Central to its parasitic lifestyle is the formation of a specialized compartment, the parasitophorous vacuole (PV), which is the parasite’s replication niche. The PV membrane (PVM), initially derived from the host cell’s plasma membrane, surrounds the replicating parasite and acts as a protective barrier against host immune mechanisms but also as a barrier for the exchange of nutrients and waste products between the parasite and the host (3). Classical transporters have not been identified within the PVM (4), instead nutrient pores formed by GRA17 and GRA23 are instrumental in modulating the selective permeability of the PVM allowing the passage of molecules up to 1300-1900 Da (5)(6).

To uncover additional genes encoding for proteins involved in nutrient acquisition from the host at the PVM, or the correct trafficking to and insertion into the PVM, in the absence of *GRA17*, we recently conducted a genome-wide synthetic lethality screen within a parasite background lacking *GRA17* (*7*). This screen discovered multiple GRA proteins (GRA57, GRA70, GRA71, GRA72) that are required for the correct localization of GRA17/GRA23 to the PVM (7,8) while other studies have identified that GRA42, GRA43, and the chaperone-like GRA45 protein are also involved in this process (8,9). *GRA72* was confirmed to be synthetically lethal with *GRA17* and deletion of *GRA72* in a wild-type background resulted in the formation of ‘bubble’ vacuoles with reduced permeability similar to *GRA17* knockout parasites (7). AlphaFold2-multimer (10,11) (12) predicted that GRA72 can form a hexameric pore (7). Because deletion of *GRA72* also led to mislocalization of GRA17/GRA23, it was unclear if the phenotype of *GRA72* knockout parasites was due to mislocalization of GRA17/GRA23 or from GRA72 itself possessing a crucial role as a pore-forming protein.

In this study we set out to better understand the functional role of GRA72 at the PVM. We show that GRA47 immunoprecipitated with GRA72 suggesting they are protein-interacting partners. Consistent with the AlphaFold prediction of a heptameric and hexameric pore for GRA47 and GRA72, respectively, blue native gel electrophoresis showed that GRA47 and GRA72 are present in large complexes. AlphaFold predictions also indicated the presence of a conserved pore-lining histidine residue in GRA47 and GRA72 pores, which we here show to play a crucial role in their function. Moreover, expression of GRA47 and GRA72 in *Xenopus* oocytes for electrophysiological analysis confirmed that GRA47 and GRA72 can modify membrane conductance and control resting membrane potential, thus further providing evidence for their role as pore-forming entities. Further investigation is warranted into the molecular mechanisms governing the permeability of the PVM and the specific molecules that pass through these pores. Overall, our results highlight the important role of *Toxoplasma* dense granule proteins GRA47 and GRA72 in modulating small molecule permeability of the PVM.

## Materials and Methods

### Site-directed mutagenesis

To mutate the conserved histidine residue of GRA47 and GRA72 to H169E, H169G, H169R, and H168E, H168G, and H168R, respectively, we employed a three-step site-directed mutagenesis protocol. Firstly, we designed primers (Table S1) targeting the desired histidine residues and used them to amplify the gene of interest using Q5 Hot Start High-Fidelity 2X Master Mix (New England Biolabs). After PCR amplification, the amplicon underwent Kinase, Ligase & DpnI (KLD) treatment (New England Biolabs). The KLD-treated reaction was then transformed using chemically competent *E. coli* cells, and the correct clones were selected by sequencing.

### Generation of parasite strains

To generate *GRA47* knockout parasites, we co-transfected the pU6-Universal plasmid carrying a sgRNA targeting the gene of interest (Table S1) and the BamHI-HF (New England Biolabs) linearized pTKOatt plasmid, which contains the *HXGPRT* selection cassette (13), into ME49 Luc+ *Δhxgprt* or RH Luc+ *Δhxgprt* parasites. The transfection ratio was 5:1 (sgRNAs: linearized plasmid). After transfection, the parasite strains were selected with 25 μg/ml mycophenolic acid (MPA) (Millipore 89287) and 25 μg/ml xanthine (Xan) (Millipore X3627). Following three rounds of drug selection with MPA-Xan, single knockout clones were isolated through limiting dilution and confirmed by PCR (Fig. S1).

To produce endogenously C-terminally tagged *GRA47*, a donor template was created by PCR amplification with forward primer (GRA47 F) (Table S1) amplifying the 5’ region of 40 base pairs homologous to the sequence immediately upstream of the stop codon of the GOI, in-frame with the 3xHA tag and the reverse primer (DHFR R) (Table S1) amplifying the *DHFR* cassette from the pLIC plasmid (14). The donor template consisted of a 5’ region of 40 base pairs homologous to the sequence immediately upstream of the stop codon of the GOI, in-frame with the 3xHA tag, and followed by a stop codon. Additionally, a 3’ region of 40 base pair homology was chosen from the downstream region of the GOI, beyond the CRISPR cut site. The pU6-Universal plasmid, containing an sgRNA targeting the gene of interest, and the donor template were electroporated into RH*Δku80Δhxgprt* (14) parasite background at a 5:1 ratio. After pyrimethamine selection of transfected parasites, individual clones were selected by limited dilution and confirmed by immunofluorescence assays and Western blotting (Fig. S2). Endotagging, knockout generation and complementation of GRA72 has been described previously (7).

The construct for GRA47 complementation was created using the pUPRT::DHFR-D plasmid backbone from Addgene (Cat#58528). The *DHFR* cassette was eliminated through PCR amplification using primers pUPRT::GOI GIB F and pUPRT::GOI GIB R (Table S1). The promoter region, approximately 1500 bp upstream of the start codon, and the coding sequence of *GRA47* were amplified using primers GOI F and GOI R (Table S1). These fragments were flanked with the HA epitope sequence before the stop codon. Additionally, the 3′-UTR region, about 500 bp in length, was amplified and assembled with the other two fragments using Gibson Assembly (New England Biolabs, Cat#E2611L). To complement *Δgra47* parasites with either wild-type or site-directed histidine mutant derivatives, RH-Luc+ *Δgra47* or ME49-cLuc+ *Δgra47* were co-transfected with plasmids containing sgRNAs specifically targeting the *UPRT* locus (Table S1) and EcoRV (New England Biolabs)-linearized pUPRT::GRA47HA plasmid at a ratio 1:5 of sgRNAs to linearized plasmid. To complement *Δgra72* parasites with site-directed histidine mutant derivatives, RHCas9 *Δgra72* parasites were co-transfected with plasmids containing sgRNAs specifically targeting the *UPRT* locus and SacI (New England Biolabs)-linearized pTwist-CMV:: GRA72-HA plasmid (7) at a ratio 1:5 of sgRNAs to linearized plasmid. Following the first egress after transfection, the parasites were selected with 10 μM 5-fluoro-2-deoxyuridine (FUDR) (Sigma–Aldrich, Cat#F0503) for three passages. Individual clones were isolated through a process of limited dilution and were subsequently verified using Western blotting and immunofluorescence assays (Fig. S2). The presence of histidine mutation was further verified from parasites by PCR amplification of the GRA47 or GRA72 locus and sequencing (Fig. S4).

### Immunoprecipitation

Five 150mm tissue culture dishes were used, each containing confluent monolayers of Human foreskin fibroblasts (HFFs). These dishes were infected with each parasite strain that carried the gene of interest (GOI) with a C-terminal 3xHA epitope tag at a multiplicity of infection (MOI) of 2. The parasites were collected after scraping the cultures and then centrifuged at 570 G for 7 minutes. The supernatant was discarded, and the resulting pellet was lysed using a lysis buffer consisted of 10mM HEPES (7.9 pH), 1.5mM MgCl_2,_ 10mM KCl, 0.1mM EDTA, 0.65% of NP-40, and 1X protease inhibitor cocktail for 30 minutes on ice. The lysis buffer was adjusted to a final pH of 7.9 by stabilizing with KOH. The lysed samples were then centrifuged at 18000 G for 30 minutes, and the resulting supernatant was incubated overnight at 4 °C with Pierce magnetic beads coupled with antibodies against the HA epitope (cat no: 88837). The magnetic beads were washed three times with a washing buffer, and Western blotting was performed before further processing the samples for peptide identification using LC-MS/MS mass spectrometry (Fig. S5). To identify putative protein interacting partners, the fold change in the unique peptide count of the sample relative to the control was calculated. Proteins showing high enrichment in the sample were considered as potential interacting partners.

### PVM GRA localization

HFFs were cultured in 24-well plates, and subsequently infected with various parasite strains, including wild-type, knockout, or complemented strains at an MOI of 0.5 for 24 hours. Certain parasites transiently expressed GRA17-HA, GRA17-V5, GRA23-HA, GRA23-FLAG, GRA47-V5, or GRA72-V5 while others did not. Following the incubation period, the cells were fixed using a 3% formaldehyde solution for 20 minutes. For GRA17 or GRA23 localization, the coverslips were blocked with a blocking buffer that contains 3% (w/v) BSA, 5% (v/v) goat serum and 0.1% Triton X-100 in PBS at room temperature for 1 h whereas for GRA47 and GRA72 localization the blocking buffer consisted of 3% (w/v) BSA, 5% (v/v) goat serum and 0.001% saponin in PBS. Subsequently, the cells were stained with rabbit anti-SAG1, rat anti-HA (Sigma–Aldrich, #11867431001), mouse anti-V5 (Thermo Scientific, #MA5-15253), mouse anti-FLAG (Sigma–Aldrich, #F3165) or rabbit anti-GRA23 antibodies. After an overnight incubation at 4°C with primary antibodies, the coverslips were subjected to staining using the following secondary antibodies: goat anti-rat Alexa Fluor 594 (Thermo Scientific, #A-11007), goat anti-mouse Alexa Fluor 594 (Thermo Scientific, #A11032), and goat anti-rabbit Alexa Fluor 488 (Thermo Scientific, #A11008), all of which were diluted 1:3000 in blocking buffer. For DNA staining, DAPI was diluted at 1:2000 and applied. To assess the localization of GRA17, GRA23, GRA47, and GRA72, we quantified a minimum of 50 vacuoles containing four or more parasites and GRA17/GRA23/GRA47/GRA72 were categorized as PVM localized, partially PVM localized, or PV lumen localized.

### Live cell imaging of PVM permeability

HFFs were cultured on glass-bottom dark 24-well plates (Greiner Bio-One) and subsequently infected with tachyzoites for a 24-hour period in regular growth media. Afterward, the cells were rinsed with Phosphate Buffer Saline (PBS), and the growth medium minus phenol red (GMPR) supplemented with 10μM 5(6)-Carboxy-2’,7’-dichlorofluorescein diacetate (CDCFDA) was added to the cells for a 10-minute incubation at 37°C. CDCFDA was prepared by diluting it sequentially into GMPR from a 10mM DMSO solution. The media containing the dye was then removed, and the cells were washed three times with PBS before being replenished with GMPR. Immediately after, the cells were subjected to imaging. At least 50 vacuoles per well were quantified and classified as either CDCFDA-positive or CDCFDA-negative.

### Parasites per vacuole counting

HFFs were cultured in 24-well plates with coverslips and subsequently infected with parasites at a multiplicity of infection (MOI) of 1, which were collected through syringe lysis. After a centrifugation step for 2 minutes at 162 G, the plates were incubated in a CO_2_ incubator at 37°C. Following an initial 4-hour incubation period, the coverslips were washed using PBS to eliminate extracellular parasites. Subsequently, another 24-hour incubation at 37°C took place. To fix the samples, the coverslips were treated with 3% formaldehyde for 20 minutes, then blocked for 1 hour using a blocking buffer containing 3% (w/v) BSA, 5% (v/v) goat serum, and 0.1% Triton X-100 in PBS. Mouse anti-GRA5 antibody (BioVision, #A1299) (1:500) was applied to identify the parasitophorous vacuole, while rabbit anti-SAG1 antibody (1:4000) was employed to label the parasites within the vacuole. For secondary labeling, anti-mouse Alexa Fluor (594) and anti-rabbit Alexa Fluor (488) antibodies were used at a dilution of 1:3000. Nucleic acids were stained with DAPI. For each experimental set, between 100 to 200 vacuoles were analyzed to determine the number of parasites within each vacuole.

### Cloning and preparation of cRNA for Xenopus oocyte injection

The pET-21(+) plasmid was utilized to generate constructs of GRA17, GRA15, GRA47, and GRA72 (Table S3). These constructs were placed under the control of the T7 promoter and inserted into EcoRI (New England Biolabs) and NotI (New England Biolabs) sites. The plasmid contains a translational enhancer derived from the 5’-UTR of the major beta-globin gene from *Xenopus laevis*. The signal peptide was removed from the *GRA15* and *GRA17* sequence, and a start codon was added upstream of the sequences whereas *GRA72* and *GRA47* do not have a predicted signal peptide and therefore the full-length sequence was used. To ensure expression in *Xenopus* oocytes, codon optimization of the open reading frame was performed. Prior to cRNA synthesis, the plasmids were linearized using EcoRV endonuclease (New England Biolabs). cRNA synthesis was accomplished through *in vitro* transcription using the HiScribe T7 ARCA mRNA Kit with tailing (New England BioLabs, Cat. # E2060S).

### Injection of cRNAs into Xenopus oocytes

Oocytes from female *Xenopus laevis* (Xenopus 1 Corp, Dexter, MI) were collected and digested as we described previously (15) according to the protocol approved by the Institutional Animal Care and Use Committee (IACUC) at the University of California, Davis and conforming to the National Institutes of Health Guide for the Care and Use of Laboratory Animals. Defolliculated oocytes (stage IV-V) were individually microinjected with GRA cRNAs (20-40 ng) or H_2_O (as a control) using the Nanoliter 2020 Injector system (World Precision Instruments, Sarasota, FL, USA). Following microinjection, oocytes were stored in plates containing ND96 solution (composition in mM: 96 NaCl, 2 KCl, 1 MgCl_2_, 1.8 CaCl_2_, 5 HEPES, adjusted to pH = 7.4) supplemented with streptomycin (50 µg/mL) plus penicillin (50 Units/mL). After cRNA injection, oocytes were stored at 16 °C for 48-72h before experiments.

### Electrophysiology

Electrophysiological data were collected using the Two-Electrode Voltage-Clamp technique (TEVC) as previously described (16). In brief, two pulled borosilicate glass micropipettes were filled with 3 M KCl, resulting in resistances ranging from 0.2 to 2 MΩ. The ionic current detected by the electrodes was amplified using an Oocyte Clamp Amplifier (OC-725C, Warner Instrument Corp., USA) and digitized by a data acquisition system (Digidata 1440A, Molecular Devices, USA). Data were sampled at 2 kHz, and analyzed using pClamp 10 software (Molecular Devices, USA). All recordings were performed at room temperature (20-22 °C). The recording solutions contained (in mM) 115 NaCl, 2 KCl, and 5 HEPES, pH adjusted to 7.40, unless otherwise specified. Voltage-current curves were obtained by analyzing the magnitude of activation currents evoked by depolarizing pulses (-120 mV to +70 mV, holding potential: -90 mV, duration of depolarization pulse: 2 s). To measure resting membrane potentials, oocytes were transferred from ND96 solution to the recording solution containing 1.8 mM CaCl_2_.

### Blue Native Polyacrylamide Gel Electrophoresis

HFFs infected with parasites expressing endogenously tagged HA epitopes at the C-terminus of GRA47 or GRA72 were collected before lysing out and were centrifuged at 570 G for 7 minutes. The pellet was subsequently lysed using a lysis buffer containing different concentrations of Triton-X or NP40, and the mixture was incubated at 4°C on ice for 30 minutes. The lysed samples were then centrifuged at 18,000 G for 30 minutes. The supernatant was combined with 4X Native Sample Buffer (ThermoFisher Scientific, BN2003) and G-250 sample buffer (ThermoFisher Scientific, BN2004), and the mixture was loaded onto a 4-16% (Bis-Tris) gradient blue Native PAGE gel. After electrophoresis, the protein was transferred to a methanol-activated PVDF membrane and immunoblotted with anti-HA antibodies.

### Protein structure predictions

Blast analysis and aligning orthologous proteins was performed using Praline (17). Prediction of transmembrane domains and signal peptides was carried out using Phobius (18,19). Phosphorylation sites were examined through data obtained from ToxoDB (20,21). AlphaFold structural predictions were executed as described previously (7), the GRA72 6-mer model was used as previously published. Briefly, the AlphaFold2-multimer v3 predictions, as described in (10), were processed through the ColabFold/Mmseqs2 workflow (version 1.5.2)(12). The workflow was executed on an Nvidia A5000 graphics card with key options being: “use_templates”: false, “num_relax”: 0, “msa_mode”: “mmseqs2_uniref_env”, “model_type”: “alphafold2_multimer_v3”, “num_models”: 5, “num_recycles”: 3, rank_by: ”multimer”. All models were depicted using UCSF ChimeraX using the rank 1 model (22).

### In vivo infection

*Toxoplasma* tachyzoites from WT (ME49 Luc+ *Δhxgprt*), ME49 *Δgra47*, ME49 *Δgra47* + H169R, and ME49 *Δgra47* + GRA47-HA parasites, were obtained by lysing host HFFs through a 27-gauge needle. These tachyzoites were used to infect six-week-old female CD-1 mice via intraperitoneal injection, with each mouse receiving 1,000 parasites. To assess parasite viability, a plaque assay was conducted immediately after infecting the mice. The mice were then observed daily and weighed every two days for a period of 30 days. At the end of this 30-day post-infection period, the mice were sacrificed, and their brains were collected to isolate tissue cysts. The mouse brain tissue was homogenized in PBS, and a fraction (1/10th) of the homogenate was fixed with ice-cold methanol. The tissue cysts were subsequently stained using DBA-FITC (Vector Laboratories, #FL-1031-5) at a 1:500 dilution and quantified under a microscope.

### Statistical analyses

GraphPad Prism software was used to conduct statistical analyses. When comparing three or more groups, analysis of variance (ANOVA) was used. In cases where there were three or more groups and a single independent variable, a one-way ANOVA with Tukey’s multiple comparisons test was used. A two-way ANOVA with Dunnett’s multiple comparison test was used to compare three or more groups involving two independent variables. When assessing the significance between two groups, a t-test was used. A statistical significance level of *P* < 0.05 was considered as indicative of significance. The presented data are represented as the mean ± standard deviation. All the data shown are derived from three or more independent experiments (except brains were isolated from WT infected mice (n = 2)), with the specific ‘n’ values provided in each figure legend. The Log-rank (Mantel–Cox) test was used to determine significance in virulence in the mouse survival experiment.

## Results

### GRA47 localizes to the parasitophorous vacuole membrane (PVM) and interacts with GRA72

To identify putative protein interacting partners of GRA72, we immunoprecipitated GRA72 from HFFs infected for 24 hours with a hemagglutinin (HA)-tagged GRA72 strain. As a control, we performed immunoprecipitation with GRA57-HA. The precipitated proteins were identified through mass spectrometry (Table 1) and ranked based on the ratio of unique peptide counts between GRA72 and the control. GRA72 emerged as the most enriched protein as expected (Table 1). GRA47 (23) also exhibited high enrichment with a unique peptide count of 10 in comparison to the control, which showed no enrichment. This suggests a direct or indirect interaction between GRA47 and GRA72. Given these immunoprecipitation findings, we decided to focus on characterizing GRA47. We endogenously tagged GRA47 with the HA-epitope at the C-terminus (Fig. S2). Within intracellular parasites, GRA47 localized to the parasitophorous vacuole membrane (PVM) and displayed colocalization with GRA5 (Fig. 1A and Fig. S2B). However, when treated with a strong detergent like Triton X-100 (TX-100), GRA47 localized predominantly to the parasitophorous vacuole lumen (Fig. S2B). In extracellular parasites, GRA47 co-localized with GRA2 within dense granules (Fig. 1B).

**FIG 1.**
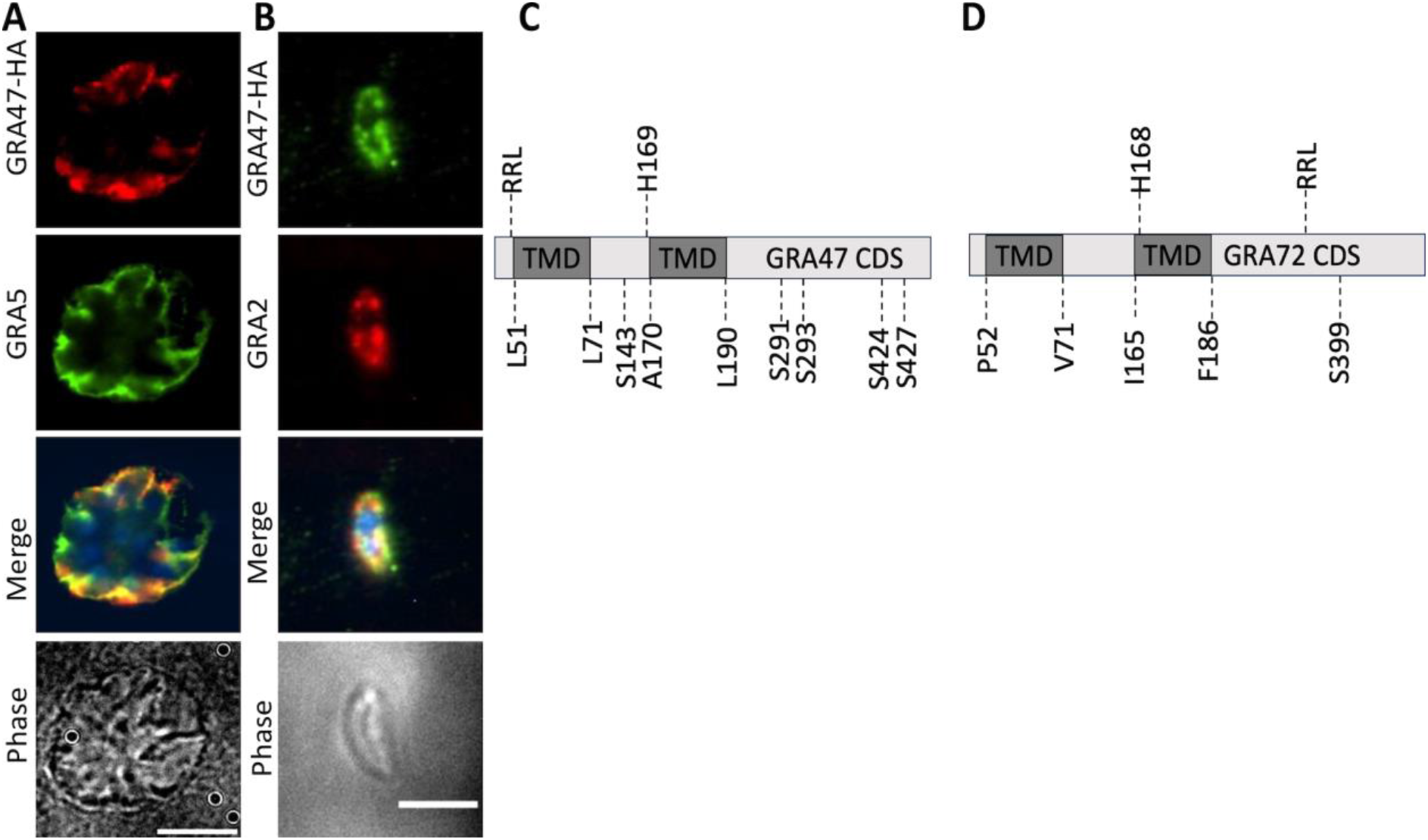
GRA47 is a dense granule protein. **A)** Immunofluorescence analysis using intracellular parasites revealed the localization of GRA47 to the parasitophorous vacuole membrane (PVM), along with co-localization with GRA5. HFFs infected with parasites for 24 hours were fixed with 3% formaldehyde and stained with anti-HA and anti-GRA5 antibodies. The scale bar used is 8 µm. **B)** Immunofluorescence analysis was performed using extracellular parasites, where GRA47 was visualized with an antibody against the HA tag co-localized with GRA2. **C and D)** Schematic diagram showing important amino acid residues of GRA47 and GRA72 (TMD= Transmembrane domain, CDS= Coding sequence, L= Leucine, A=Alanine, P= Proline, V= Valine, I=Isoleucine, F=Phenylalanine, S= Serine, RRL= *Toxoplasma* export element (TEXEL motif).

**Table 1.** Immunoprecipitation of GRA72 followed by mass spectrometry (IP-MS) analysis. Mass spectrometry was conducted on the immunoprecipitated material, and the count of unique peptides was quantified for each identified protein. The identified *Toxoplasma* proteins were then ranked based on the number of unique peptide counts in the strain expressing GRA72-3×HA compared to the control strain. This ranking was determined by calculating the fold change in relation to the control strain. If the control strain had no peptides the total number of peptides detected in GRA72-3×HA is indicated. *Toxoplasma* genes that had at least a 5-fold enrichment compared to immunoprecipitation of the control protein were considered.

| GeneID TGGT1_ | Description | Peptide fold change<br>GRA72 vs. GRA57 |
| --- | --- | --- |
| 272460 | GRA72 | 22 |
| 254000 | GRA47 | 10 |
| 202620 | GRA64 | 6 |
| 224740 | hypothetical protein | 5 |
| 230940 | hypothetical protein | 5 |

Blast analysis and subsequent alignment of orthologous proteins showed that GRA47 is highly conserved among Sarcocystidae and Eimeriidae (Supplementary data 8) but is absent in *Plasmodium* spp. and *Cryptosporidium* spp. *GRA47* is also expressed across the different *Toxoplasma* life stages (ToxoDB.org). Additionally, GRA47 is phosphorylated at multiple serine residues (Fig. 1C) some of which likely by the WNG1 kinase (24). In contrast, GRA72 orthologues were only identified in Toxoplasmatinae and only one serine residue has been shown to be phosphorylated. Both GRA47 and GRA72 have two predicted transmembrane domains (Fig. 1C and 1D) and do not have a predicted signal peptide.

### AlphaFold-multimer predictions show GRA47 and GRA72 likely form pores with a conserved pore-lining histidine

As mentioned earlier, when mild detergent solubilization such as saponin is used, GRA47 is predominantly localized to the PVM (Fig. 1A and Fig. S2B) similar to what we observed for GRA72 (7). Furthermore, both *Δgra47* and *Δgra72* parasites form bubble-like vacuoles and exhibit a reduction in CDCFDA permeability, which are typical characteristics of pore-forming proteins that we have previously studied (5). Therefore, we hypothesized that both GRA47 and GRA72 form a pore in the PVM that is essential for the flow of small molecules. To test this hypothesis, we recently utilized AlphaFold predictions on GRA72 and showed that AlphaFold-multimer (10) predicts GRA72 to form hexameric pore-like structures (7). We applied a similar AlphaFold model and found that AlphaFold predictions for the helical domain of GRA47 reveals a potential pore-forming complex when modeled as a heptamer (Fig. 2A and 2B). While the initial N-terminal domain (residues 1-159) forms a domain with many random coil regions, the rest of the C-terminus folds as a structured pore through a long α-helix (residues 162-235) which forms a funnel reminiscent of the *Plasmodium* Exp2 heptamer found within the *Plasmodium* PTEX cryo-EM structure (25) in its engaged state (Fig. S3A). Within this structured core, the predicted local distance difference test (pLDDT) score is high (Fig. 2A), suggesting a good confidence, and the pore also forms comparatively when the input prediction is truncated to start at residue 161 (Fig. 2A). Furthermore, pore formation is also observed within the closest orthologs found in related apicomplexan species *Neospora caninum*, *Eimeria tenella*, *Sarcocystis neurona*, and *Besnoitia besnoiti* (Fig. S3B). The charge potential of the pore is noteworthy. When placed on a phospholipid bilayer, both upper and lower entry points display a positively charged surface, as predicted by coulombic charge calculations (Fig. 2B). One of the constriction points of the pore channel, where the main α-helix crosses over in between monomers, is defined by a histidine (residue 169) (Fig. 2C). Close to this histidine is another histidine and a phenylalanine, which form other constriction points in the pore. Similar basic residues together with charged residues are also observed at constriction points of other pore-forming predictions of 7-mer GRA17 (Fig. 2F), 6-mer GRA72 (Fig. 2D), and within the experimental 7-mer *Pf*Exp2 structure (Fig. 2E). These gating residues all seem to constrict the pore channel within a range of 15 to 18Å.

**FIG 2.**
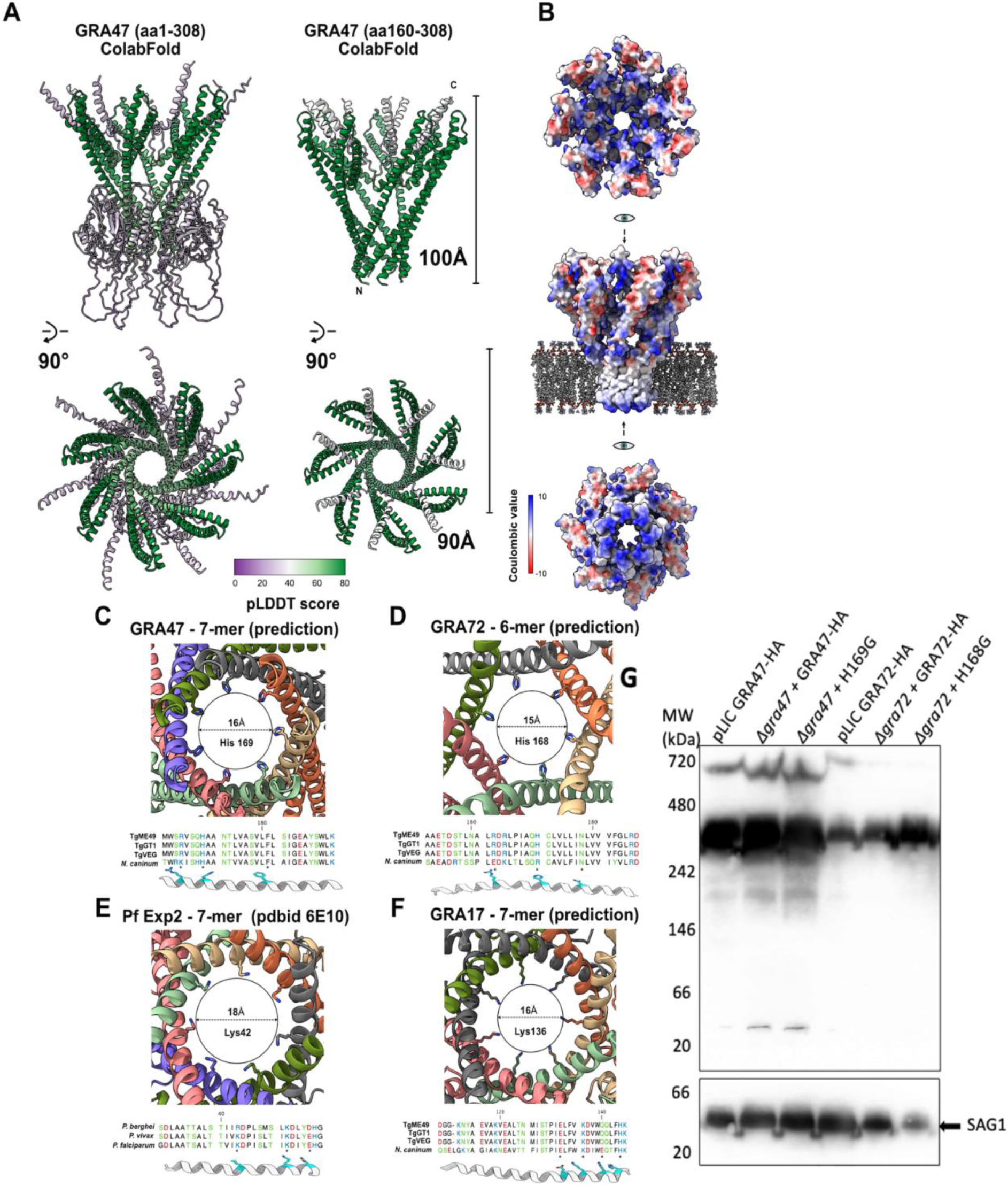
AlphaFold-multimer predicts GRA47 to form a pore. **A)** Overall successful pseudosymmetry assembly of GRA47 in both full length and truncated form. Structures are depicted in a cartoon diagram fashion and colored using the calculated pLDDT score. **B)** Using the coulombic surface charge, GRA47 is represented in a top, side and bottom view. Phospholipid bilayer was added to the side view for scale and docked manually against the lower pore funnel. **C-F)** Conserved basic residues line the bottom pore constriction of GRA47 (C), GRA72 (D), PfExp2 (E) and GRA17 (F). Using always the same rules of representation, the tightest pore passage is depicted in a cut slice highlighting the residue gating the pore passage. Every monomer is colored separately while the pore limits are schematically shown as a circle with the diameter measured. For every example, a lower sequence alignment is shown with the corresponding transmembrane a-helix from which residues closing off the pore are colored in cyan. In the case of PfExp2, sequence comparison between *P. falciparum*, *P. vivax* and *P. berghei* while for GRA47/GRA72/GRA17, sequence comparisons are made between TGME49, TGGT1, TGVEG and *N. caninum.* **H)** A pellet harvested from HFFs infected with the indicated parasites before lysis was subjected to extraction using 1% TX-100. The resulting samples were then centrifuged, and the supernatants were utilized for blue native polyacrylamide gel electrophoresis (BN-PAGE). Following BN-PAGE, Western blotting was performed sequentially using anti-HA and anti-SAG1 antibodies.

As mentioned before, Alphafold-multimer predictions indicated that both GRA47 and GRA72 can form multimers, leading to the formation of pores. This prediction aligns with the fact that both proteins possess at least one transmembrane domain, allowing them to assemble and form complexes capable of creating large pores in phospholipid membranes. To confirm this prediction, we conducted blue native polyacrylamide gel electrophoresis (BN-PAGE) to determine whether GRA47 and GRA72 form multimers. We collected lysate from HFFs infected with parasites expressing HA-tagged GRA47 or GRA72 before the parasites lysed out and extracted them using two different TX-100 concentrations or NP-40, subjecting the soluble extracts to BN-PAGE and Western blot analysis (Fig. 2G, Fig. S6). At 1% TX-100, both GRA47-HA and GRA72-HA formed complexes within the size range of 242-480 kDa, while 0.5% TX-100 and 0.65% NP-40 were not able to efficiently extract these proteins from the pellet (Fig. 2G, Fig. S6). A small fraction of GRA47-HA was also detected in its monomeric form (Fig. 2G). To determine if the pore-lining histidine mutation in GRA47 or GRA72 affected complex formation, we performed a similar blue native polyacrylamide gel electrophoresis on HFFs infected with GRA47 H169G or GRA72 H168G. H169G and H168G proteins migrated similarly to wild-type proteins (Fig. 2G), indicating that mutation of the pore-lining histidine likely did not alter formation of a larger complex. In conclusion, AlphaFold predictions corroborate the biochemical evidence that GRA47 and GRA72 likely form multimeric pores on the PVM, each featuring a conserved pore-lining histidine, thereby underscoring their possible functional similarities.

### Deletion of GRA47 leads to formation of bubble vacuoles and reduced PVM small molecule permeability

To conduct a functional analysis of GRA47, we generated a separate knockout for this purpose (Fig. S1). Additionally, we performed a complementation experiment by introducing a C-terminally HA-tagged version of GRA47 into the *Δgra47* parasites (Fig. S2). Our observations revealed that the *Δgra47* parasites formed significantly enlarged ‘bubble’ vacuoles with irregular morphology (Fig. 3A). Furthermore, some of these vacuoles appeared to have collapsed with opaque parasites inside, similar to other bubble vacuole forming parasites we characterized previously (5,7). In our previous studies, we demonstrated that parasite strains that form bubble vacuoles also have reduced permeability to small molecules (5,7). To determine whether *Δgra47* parasites have decreased permeability to small molecules, we utilized the properties of the small (445.2 Da) vital dye 5-(and-6)-Carboxy-2′,7′-Dichlorofluorescein Diacetate (CDCFDA). CDCFDA can pass through cell membranes and remains non-fluorescent until it enters living cells, where intracellular esterases convert it into the fluorescent form, which cannot pass through cell membranes. CDCFDA can enter the PV through passive diffusion, as it is smaller than the established size exclusion limit of the *Toxoplasma* PVM (5)(6). PVs generated by *Δgra47* parasite had significantly reduced permeability to the dye, with only about 40% of vacuoles showing permeability (Fig. 3B and 3C). As seen previously, only 20% of PVs generated by *Δgra17* parasites were permeable to the dye (Fig. 3B and 3C). When we reintroduced a C-terminally HA-tagged version of *GRA47* into the *Δgra47* parasites (Fig. S2), the PV permeability to the dye and the wild-type vacuole phenotype were restored (Fig. 3A).

**FIG 3.**
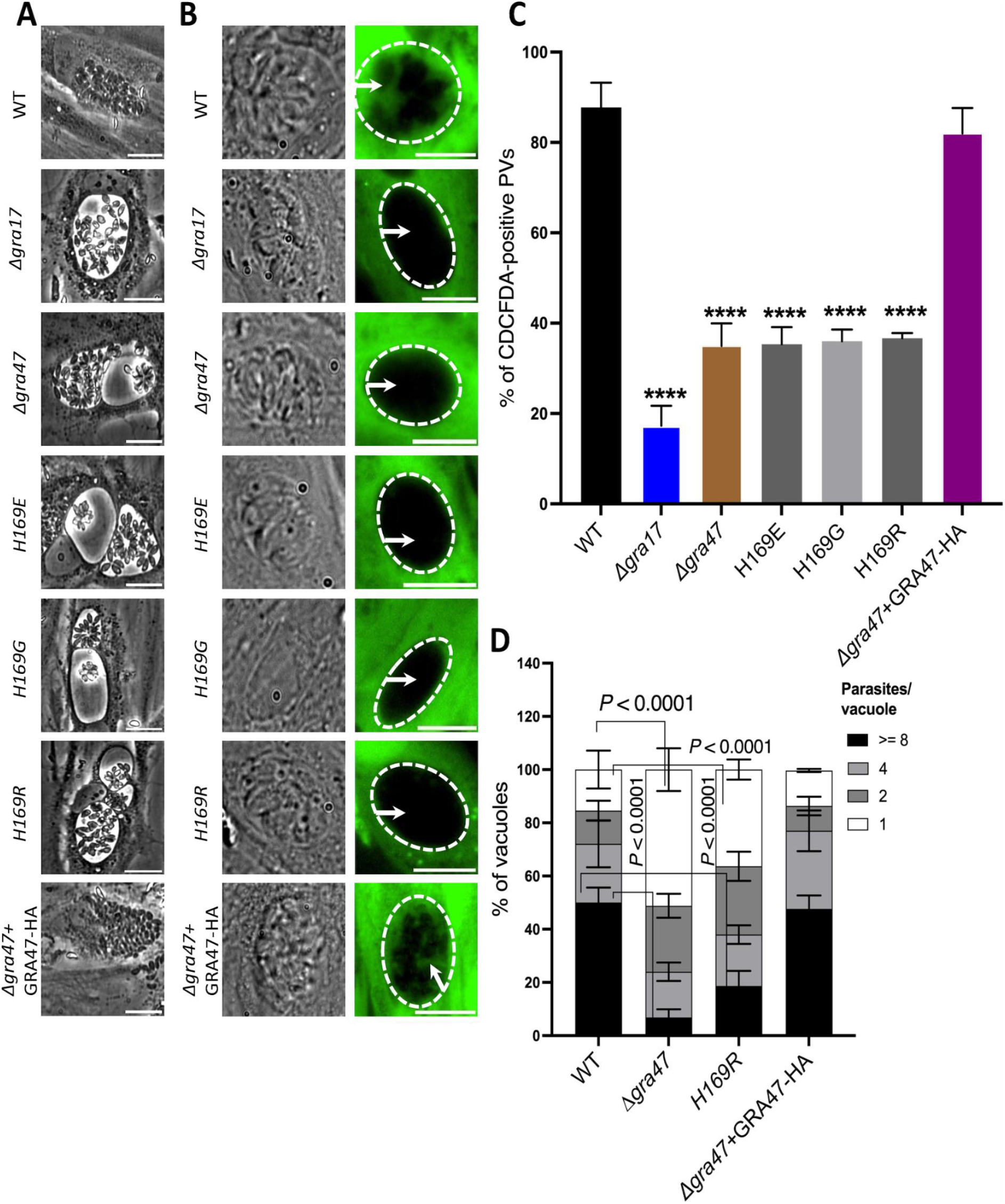
The GRA47 pore-lining histidine is important for its function. **A)** Human foreskin fibroblasts (HFFs) were infected with the specified strains at an MOI of 1 for 36 hours (WT= wild-type). Live imaging was carried out using phase contrast microscopy, and the resulting images were captured. The scale bar in the images corresponds to 80 µm. **B)** HFFs were infected with the designated parasite strains for 24 hours. Following this, they were exposed to CDCFDA (5-(and-6)-Carboxy-2’,7’-dichlorofluorescein diacetate) for 10 minutes. Subsequently, the dye was washed away using PBS and imaging was performed. Shown are representative images, with the wild-type, knockout and complemented parasite strains. Scale bar corresponds to 75 µm. **C)** The percentage of CDCFDA-fluorescent vacuoles was quantified for each strain. A minimum of 50 vacuoles per well were assessed and categorized as either CDCFDA-positive or CDCFDA-negative. The results are shown as mean ± SD from three independent experiments. One-way ANOVA with Tukey’s multiple comparison test was used to determine significance (\*\*\*\**P* < .0001, for WT, *Δgra17, Δgra47 and Δgra47+*GRA47-HA, n=6, for H169E, H169G and H169R, n=3.). **D)** HFFs were plated in 24-well plates with coverslips and subsequently infected with various strains of parasites at a multiplicity of infection (MOI) of 1 for 24 hours. After the infection period, the cells were fixed using a 3% paraformaldehyde solution and subjected to blocking using a blocking buffer. The coverslips were then stained with rabbit anti-SAG1 antibody. In each experiment, a total of 100 to 200 vacuoles were analyzed. The data are presented as the average values along with the standard deviation (±SD). To analyze the results, a two-way analysis of variance (ANOVA) was performed, followed by Tukey’s multiple comparison test (for WT and *Δgra47*, n=5, for H169R and *Δgra47+*GRA47-HA, n=3.

AlphaFold predicted that the lumen of the pore formed by GRA47 and GRA72 is lined with histidine residues (Fig. 2G), which might regulate the movement, flow, or transfer of small molecules. To investigate the potential role of histidine residues in pore function, we conducted mutations, replacing histidine with either a negatively charged (H->E), small (H->G), or positively charged (H->R) residue. After creating these mutations, we complemented *Δgra47* knockout parasites with plasmids encoding the wild-type or the histidine mutant versions of GRA47 (Fig. S2). Complementation of *Δgra47* parasites with GRA47 containing histidine substitutions failed to restore the wild-type vacuole phenotype (Fig. 3A). Additionally, PVs formed by these GRA47 histidine mutants (H169E, H169G, and H169R) had reduced dye permeability (Fig. 3B and 3C), with fewer than 40% of vacuoles showing any permeability to the dye. In contrast, over 80% of vacuoles formed by *Δgra47* parasites complemented with wild-type GRA47 demonstrated dye permeability (Fig. 3B and 3C). Additionally, we observed a significantly lower percentage of *Δgra47* vacuoles containing 8 parasites and a significantly higher percentage of vacuoles containing 1 parasite, compared to the vacuoles of wild-type parasites, indicating that the growth rate of *Δgra47* parasites is slower (Fig. 3D). This is consistent with the CRISPR fitness score of -3.49 (ToxoDB.org) reported for this gene (26). Complementation of *Δgra47* parasites with a wild-type copy of *GRA47* rescued this growth phenotype while complementation with the GRA47 H169R mutation did not. Overall, these data show that the deletion of *GRA47* results in aberrant ‘bubble’ vacuoles and compromised PVM permeability to small molecules, an effect reversible by wild-type *GRA47* complementation but not by histidine-substituted variants, implicating the histidine residue in pore functionality.

### The pore-lining histidine is necessary for the functioning of GRA72 pores

To investigate the potential role of histidine residues in GRA72 pore function, we performed similar histidine replacements as described for GRA47 and complemented *Δgra72* parasites with plasmids encoding the wild-type or the histidine mutant versions GRA72 (Fig. S2). *Δgra72* parasites complemented with histidine mutants continued to exhibit the formation of bubble vacuoles (Fig. 4A).

Moreover, only 35% of *Δgra72* parasites PVs complemented with GRA72 histidine changes (H168E, H168G, and H168R) were permeable to CDCFDA, which was a significant reduction compared to *Δgra72* parasites complemented with wild-type GRA72 or wild-type parasites (Fig. 4B and 4C). As previously reported, *Δgra72* parasites grew slower compared to wild-type parasites. Complementation with a wild-type copy of GRA72 rescued this growth phenotype while complementation with the GRA72 H168R mutation did not (Fig. 4D). In conclusion, these findings underscore the importance of the histidine residues in the functioning of GRA72 pores.

**FIG 4.**
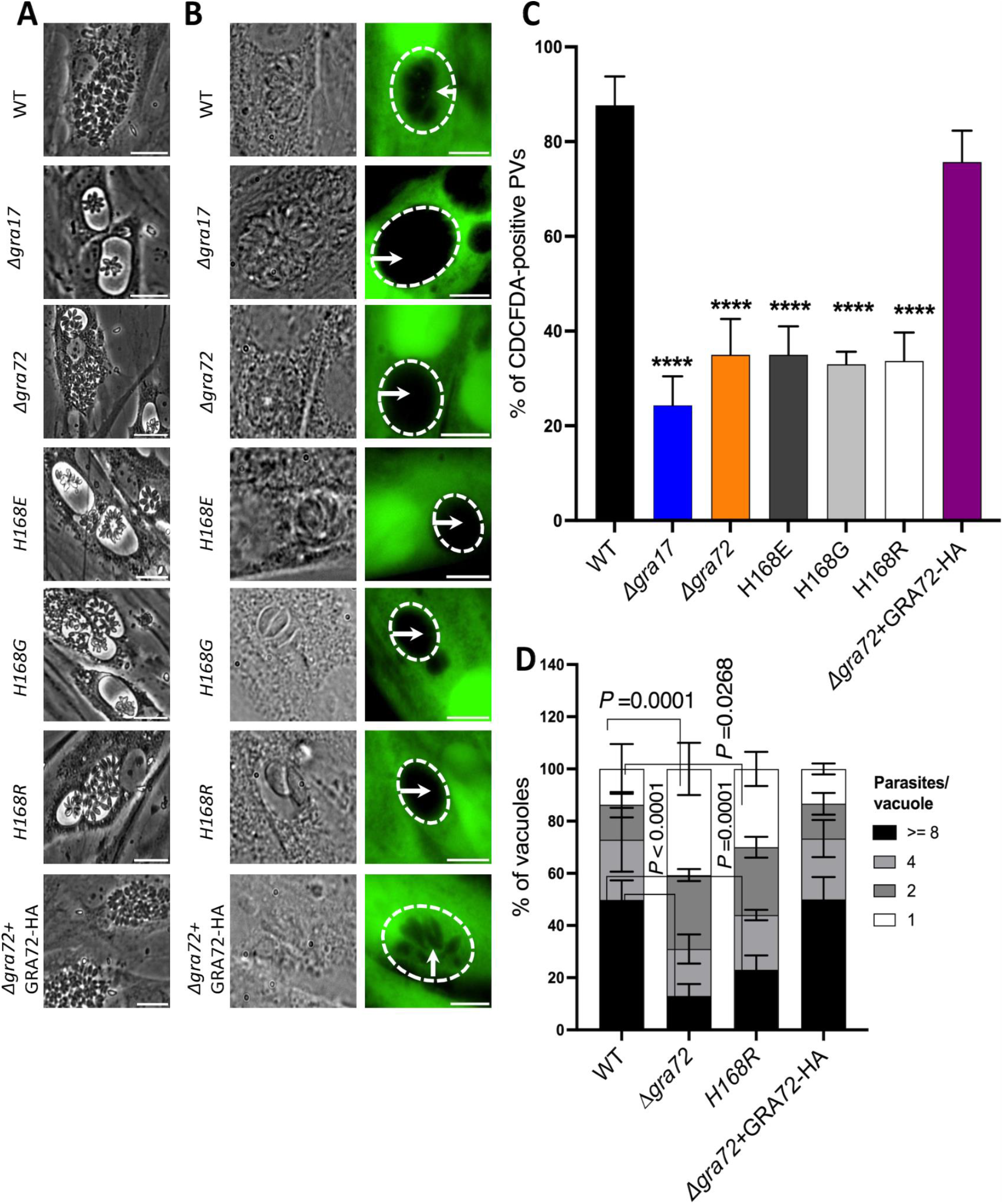
The GRA72 pore-lining histidine is important for its function. **A)** Human foreskin fibroblasts (HFFs) were infected with the specified strains at an MOI of 1 for 36 hours (WT= wild-type). Live imaging was carried out using phase contrast microscopy, and the resulting images were captured. The scale bar in the images corresponds to 80 µm. **B)** HFFs were infected with the designated parasite strains for 24 hours. Following this, they were exposed to CDCFDA (5-(and-6)-Carboxy-2’,7’-dichlorofluorescein diacetate) for 10 minutes. Subsequently, the dye was washed away using PBS and imaging was performed. Shown are representative images of the wild-type, knockout and wild-type or Histidine-mutant complemented parasite strains. Scale bar used is 75 µm. **C)** The percentage of CDCFDA-fluorescent vacuoles was quantified for each strain. A minimum of 50 vacuoles per well were assessed and categorized as either CDCFDA-positive or CDCFDA-negative. The results are shown as mean ± SD from three independent experiments. One-way ANOVA with Tukey’s multiple comparison test was used to determine significance (\*\*\*\**P* < .0001, n = 3). **D)** HFFs were plated in 24-well plates with coverslips and subsequently infected with various strains of parasites at a multiplicity of infection (MOI) of 1 for 24 hours. After the infection period, the cells were fixed using a 3% paraformaldehyde solution and subjected to blocking using a blocking buffer. The coverslips were then stained with rabbit anti-SAG1 antibody. In each experiment, a total of 100 to 200 vacuoles were analyzed. The data is presented as the average values along with the standard deviation (±SD). To analyze the results, a two-way analysis of variance (ANOVA) was performed, followed by multiple comparisons.

### The phenotype of Δgra47 parasites is not due to mislocalization of GRA17, GRA23, or GRA72 to the PVM

Because deletion of *GRA17*, *GRA23*, or *GRA72* can lead to the formation of ‘bubble’ vacuoles with reduced PVM permeability (5,7) it was possible that the phenotype of Δ*gra47* parasites was due to mislocalization of these proteins. To investigate whether the proper localization of these dense granule proteins is affected in Δ*gra47* parasites, we ectopically expressed GRA17, GRA23, and GRA72 and then quantified their localization to the PVM or the PV lumen. There was no statistical difference in PVM localization of GRA17, GRA23, or GRA72 between Δ*gra47* and wild-type parasites (Fig. 5 and Fig. 6). From these data, it can be inferred that the formation of bubble vacuoles in *Δgra47* parasites is not a result of altered localization of GRA17, GRA23, or GRA72. Parasites lacking GRA47 exhibited comparable export levels of GRA16 and GRA24, mirroring the export levels observed in wild-type parasites. This implies that the deletion of GRA47 does not impact GRA export to the host nucleus (Fig. S7). Additionally, our recent findings demonstrated that GRA72 similarly does not contribute to GRA export (7)

**FIG 5.**
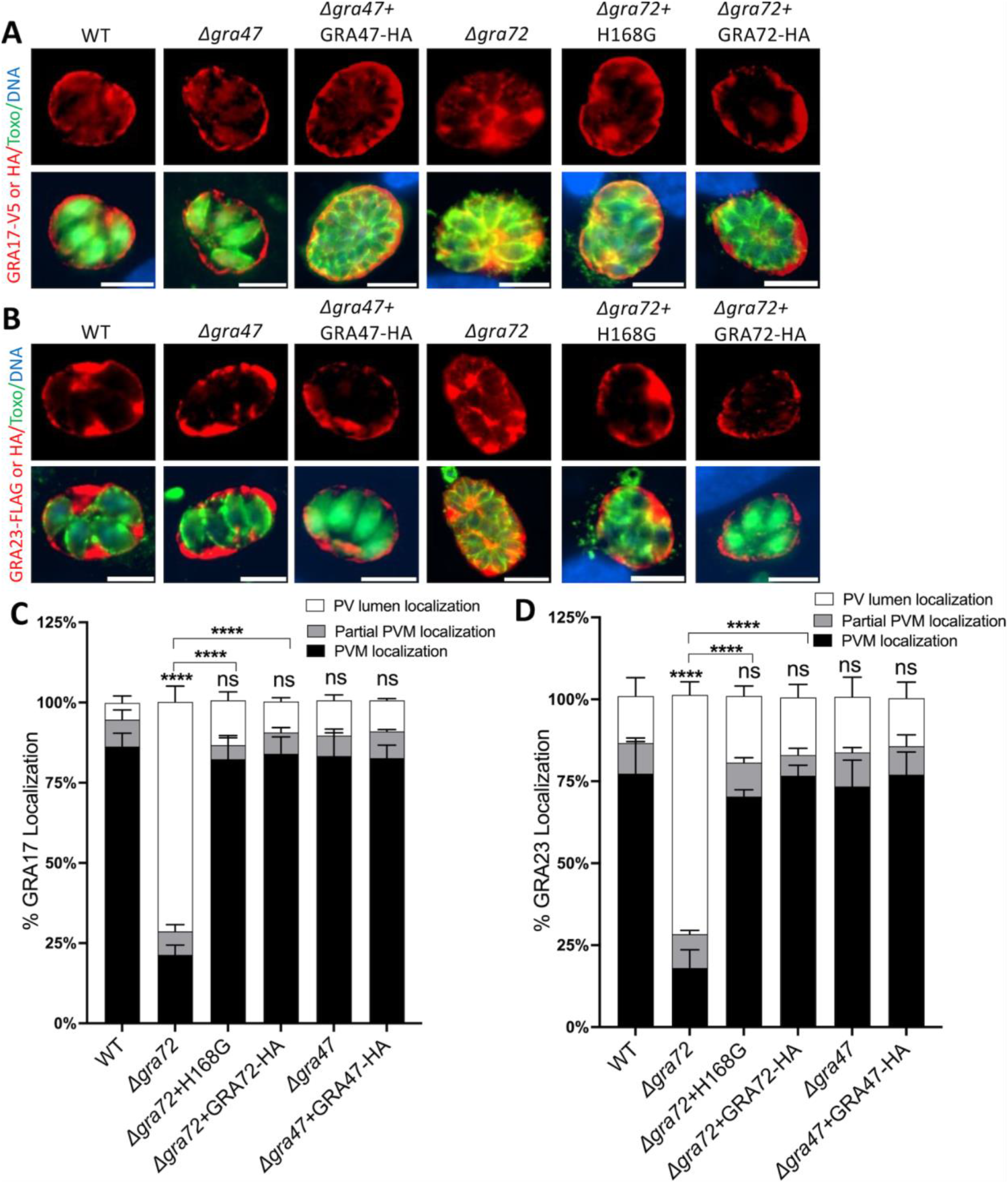
Localization of GRA17 and GRA23 in knockout parasites. HFFs were infected with the indicated parasite strains transiently expressing GRA17-HA, GRA17-V5, GRA23-HA, or GRA23-FLAG at an MOI of 0.5 for 24 h. Shown are representative images of **A)** GRA17, **B)** GRA23.The scale bar used was 8μm. **C)** The percentage of vacuoles with PVM **C)** GRA17 **D)** GRA23 was quantified. Two-way ANOVA Dunnett’s multiple comparisons test was used to establish the significance of the observed results (\*\*\*\**P* < .0001 n = 3 ns= non-significant).

**FIG 6.**
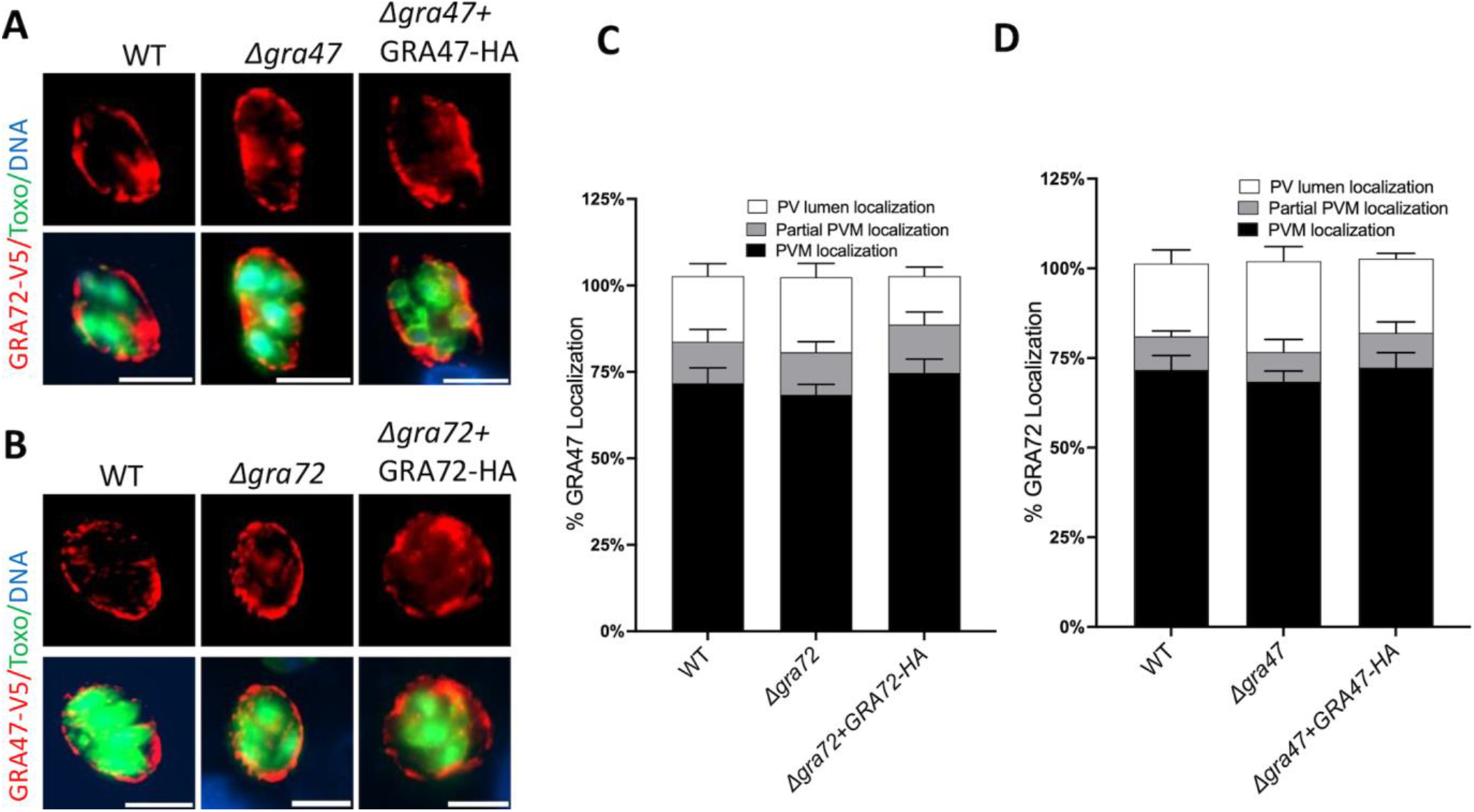
Localization of GRA47 and GRA72 in knockout parasites. **A)** HFFs grown in 24 well plates were infected with the indicated parasite transiently expressing GRA47-V5 or GRA72-V5 at an MOI of 0.5 for 24 h. Shown are representative images of **A)** GRA72, **B)** GRA47. The scale bar used was 8μm. **C & D**) The percentage of vacuoles with PVM, partial or PV Lumen GRA47 or GRA72 was quantified (n = 3). Two-way ANOVA Dunnett’s multiple comparisons test was used to establish the significance of the observed results.

### The phenotype of Δgra72 parasites is not due to mislocalization of GRA17, GRA23, or GRA47 to the PVM

Previously, it was demonstrated that deletion of *GRA72* also led to mislocalization of GRA17/GRA23 (7,27). It was therefore unclear if the phenotype of *GRA72* knockout parasites was due to mislocalization of GRA17/GRA23, or potentially of GRA47, or from an intrinsic pore-forming activity of GRA72. To determine this, we investigated whether the localization of GRA17 and GRA23 was affected in parasites with a histidine mutation in GRA72. *Δgra72* parasites complemented with GRA72 containing a mutation of the GRA72 histidine to glycine had normal localization of GRA17 and GRA23 (Fig. 5). Thus, although the GRA72 histidine is important for GRA72’s role in maintaining normal PV morphology and permeability, this histidine is not important for GRA72’s role in the correct localization of GRA17 and GRA23. We also tested if the localization of GRA47 was affected in *Δgra72* parasites and observed that GRA47 localization was normal (Fig. 6). Therefore, the phenotype observed in *Δgra72* parasites is unlikely due to the mislocalization of GRA17, GRA23, or GRA47.

### GRA47 and GRA72 alter the ionic currents and resting membrane potential of ***Xenopus* Oocytes**

To confirm the role of GRA47 and GRA72 as pore-forming proteins, we utilized *Xenopus laevis* oocytes as a heterologous system, similar to our previous demonstration of GRA17 pore formation (5). We employed *in vitro* transcribed cRNAs derived from plasmids containing the genes for GRA47 or GRA72, that were injected in *Xenopus* oocytes to assess their potential to control the membrane conductance. We included GRA15, a non-channel protein of *Toxoplasma* known to be non-functional in this context, as the negative control, and GRA17, previously shown to alter membrane conductance and form a pore (5), as the positive control. The electrophysiological data revealed that injecting GRA17, GRA47 and GRA72 in cRNAs into *Xenopus* oocytes led to a slight depolarization of the resting membrane potential (Fig. 7A). Consistently, oocytes injected with GRA17, GRA47 and GRA72 exhibited notably higher ionic currents than those injected with water or GRA15 (Fig. 7C and 7D). The relationship between current and voltage showed an increased linear correlation in the net membrane current (Fig. 7C). Collectively, these data align with our hypothesis that exogenously expressing GRA47 or GRA72 in cell membranes can lead to the formation of pores.

**FIG 7.**
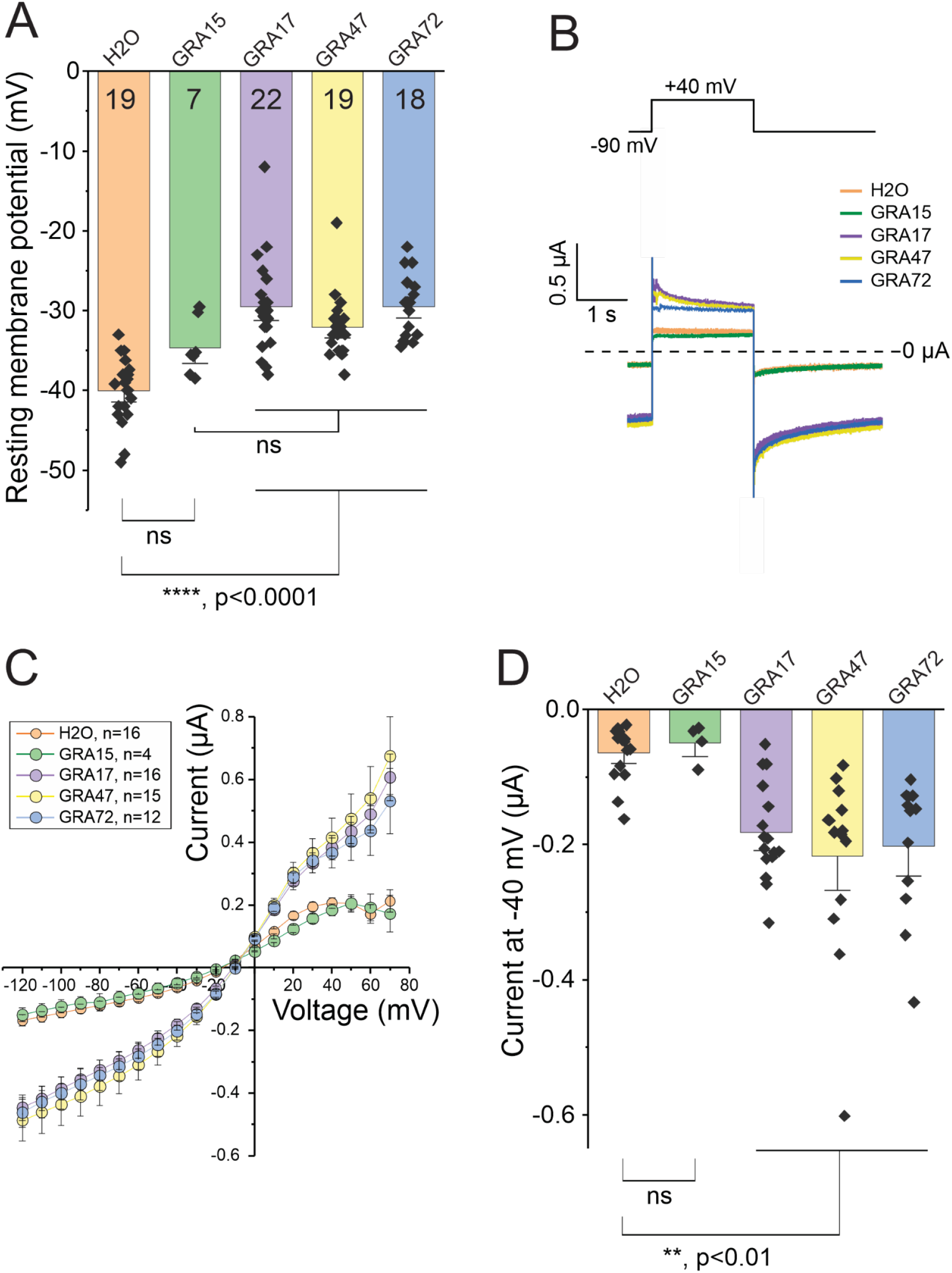
GRA47 and GRA72 change the membrane conductance characteristics of *Xenopus* oocytes. **A)** Average resting membrane potential (RMPs) of injected oocytes. Statistics: one-way ANOVA followed by Tukey’s Multiple comparisons. **B)** TEVC protocol and representative ionic current traces recorded in oocytes expressing GRA15, GRA17, GRA47, or GRA72. **C)** I-V relationship from data shown in B. **D)** Quantification of current recorded at holding potential -40mV. Statistics: one-way ANOVA followed by Tukey’s Multiple comparisons.

### GRA47 is required for the in vivo proliferation and pathogenicity of Toxoplasma

To assess the significance of GRA47 in the *in vivo* proliferation and pathogenicity of *Toxoplasma,* we conducted intraperitoneal (i.p.) infections in CD-1 mice, using 1,000 tachyzoites from different strains: WT (ME49 Luc+ *Δhxgprt*), ME49 *Δgra47*, ME49 *Δgra47* + H169R, and ME49 *Δgra47* + GRA47-HA parasites. Unlike mice infected with Δ*gra47* and ME49 Δ*gra47* + H169R parasites, those infected with wild-type parasites experienced a reduction in body weight (Fig. 8A) and a decline in overall body condition throughout the course of the experiment. Among the mice infected with wild-type parasites, only two out of five survived (Fig. 8B), while four out of five mice infected with *Δgra47* parasites survived for the duration of the experiment.

**FIG 8.**
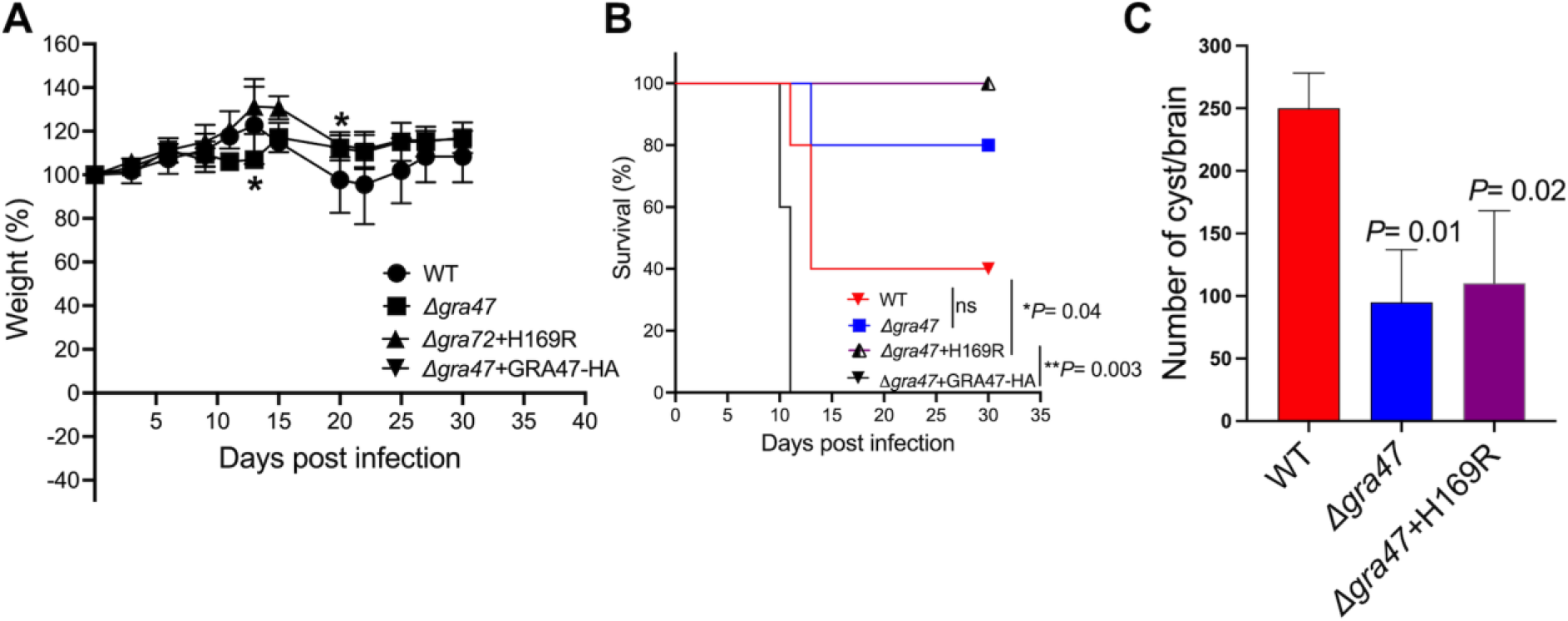
*Δgra47* and H169R parasites are less virulent in mice. **A)** CD1 mice (n=5) were infected intraperitoneally with 1,000 tachyzoites of WT, *Δgra47*, *Δgra47* + H169R, or *Δgra47* + GRA47-HA, all belonging to the ME49 type II strain. Over the course of the infection period, the mice were consistently monitored and weighed regularly. The weight data were then plotted as the average change in body weight for each group, with the weight on the day prior to infection set as 100%. To assess the statistical significance, a one-way ANOVA with Dunnett’s multiple comparison test was used (*\* P*= 0.04). **B)** The survival of mice after infection was monitored regularly for 30 days. Statistical significance was determined using the Log-rank (Mantel–Cox) test. **C)** Surviving CD-1 mice were sacrificed 30 days post infection and the number of cysts/brain was quantified. Brains were isolated from WT (n = 2), *Δgra47* (n = 4), and *Δgra47* + H169R (n=5) infected mice and cyst numbers were quantified. One way-ANOVA with Dunnett’s multiple comparisons test was used to determine statistical significance and error bars indicate SD.

Conversely, none of the mice infected with ME49 *Δgra47* + GRA47-HA parasites survived throughout the experimental period (Fig. 8B). Notably, all mice infected with ME49 *Δgra47* + H169R parasites survived the full duration of the experiment. Mice infected with *Δgra47* parasites had a lower cyst burden (averaging 95 cysts), while the surviving mouse infected with wild-type parasites had an average of 250 cysts in the brain (Fig. 8C). Similarly, mice infected with ME49 *Δgra47* + H169R parasites also exhibited a reduced cyst count in the brain (averaging 110 cysts) compared to the wild-type (Fig. 8C). These findings collectively indicate that GRA47 is important for the *in vivo* virulence and proliferation of *Toxoplasma*.

## Discussion

The interaction between the parasite and its host has long been recognized as fundamental to *Toxoplasma*’s successful establishment within its host. However, the specific mechanisms by which it acquires host-derived nutrients are incompletely understood. The findings presented in this study contribute to understanding of some of these mechanisms by highlighting the role of GRA47 and GRA72, two *Toxoplasma* dense granule proteins, as pore-forming proteins that influence small molecular permeability across the PVM.

The ‘bubble vacuole’ phenotype in parasites lacking pore-forming proteins -GRA17, GRA23, GRA72, and GRA47-suggests a perturbation in molecular transport, ion homeostasis, and/or osmotic regulation within the PV, thereby affecting vacuolar morphology. The observed decreased dye permeability across the PVM in these knockout strains also suggests compromised molecular trafficking. The shared phenotype among parasites lacking different pore-forming proteins raises the question if these proteins operate redundantly or synergistically. The partial retention of dye permeability, even in the absence of one of these key pore-forming proteins, points to functional redundancy. However, using live imaging, we have previously observed that a significant fraction of *Δgra17* bubble vacuoles collapse, and it is likely that the PVM in these collapsed PVs is broken. It therefore remains ambiguous if the fraction of vacuoles of these GRA knockout strains that are permeable to the dye are indicative of collapsed vacuoles or a true indicator of functional pores. To resolve this a more careful examination of the phenotype is required: microinjection of different dyes, of smaller and larger size than the pore exclusion limit, could be used to discern collapsed vacuoles (permeable to both small and large dyes) from truly permeable ones (permeable only to small dyes). Furthermore, generating combinatorial knockout strains could help elucidate potential functional redundancies or interdependencies. Although this might be complicated by the synthetic lethality observed in some pairs (e.g., *GRA23*-*GRA17*, *GRA72*-*GRA17*). Another avenue is to assess whether overexpression of one pore-forming protein can restore the wild-type phenotype in the absence of another, similar to our previous work demonstration that overexpression of GRA23 or *Plasmodium falciparum* EXP2 in a *Δgra17* background reverts phenotypes to wild-type. Even when these proteins form distinct pores, the absence of one could still attenuate the functionality of the remaining pores, thereby influencing the observed phenotype. This scenario is conceptually similar to symport and antiport systems, in which the directional flux of one ion or molecule can modulate the transport kinetics of another molecule, either in the same or opposite direction. Thus, the pores may not operate in isolation, but could be part of an interconnected system that regulates molecular and ionic flux across the PVM.

Our results indicate two distinct roles for GRA72: 1) regulating the trafficking or anchoring of GRA17 and GRA23 within the PVM (7,27), possibly through protein-protein interactions or complex assembly; 2) serving as a pore within the PVM. These roles may not be mutually exclusive and could be functionally intertwined. For instance, GRA72 may impact the lipid and cholesterol composition of the PVM, thereby modulating the formation of microdomains, which in turn could affect the localization of proteins like GRA17 and GRA23. Our data show that the localization of GRA17 and GRA23 is normal in strains with mutations to the pore-lining histidine residue in GRA72, even though they compromise the normal shape of the PV and its permeability. This suggests that the histidine residue’s primary function is related to the pore activity of GRA72, rather than the anchoring or localization of GRA17 and GRA23 within the PVM.

The conserved nature of GRA47 suggests that it serves essential functions across various coccidian species, possibly in processes related to permeability or molecular transport, given its role in *Toxoplasma*. The preservation of specific structural elements, such as the histidine residue, implies its functional importance, possibly in ion coordination or other crucial aspects of pore activity. On the other hand, the lack of conservation in GRA72 suggests unique adaptations or functionalities that are specific to *Toxoplasma* or closely related parasites. GRA47, as a conserved protein, might represent a potential target for broad-spectrum interventions against coccidian parasites, considering its shared importance across species.

While the results of this study significantly advance our understanding of the role of GRA47 and GRA72 in small molecule permeability, additional research is needed to understand the selectivity of these pores and investigate their specificity for particular molecules. Additionally, the interaction between GRA47, GRA72, and other pore-forming proteins, as well as their integration into the larger nutrient acquisition strategy of *Toxoplasma*, warrants further investigation. In conclusion, this study provides compelling evidence for the involvement, either direct or indirect, of GRA47 and GRA72 in mediating small molecule permeability across the PVM of *Toxoplasma gondii*. The identification and functional characterization of these pore-forming proteins contribute to our understanding of the intricate strategies employed by the parasite to manipulate host environments and acquire essential nutrients.

## Acknowledgements

This research received funding from the National Institutes of Health (NIH), specifically grants R21AI149071 and R21AI151084, which were awarded to J.P.J.S. Our gratitude goes to Dr. M. A. Hakimi for generously sharing the plasmids expressing GRA16-Ty and GRA24-Ty. We also extend our appreciation to Dr. Tatsunori Masatani for providing us with anti-GRA23 antibodies. Additionally, we would like to acknowledge the entire team at EupathDB.org; for their invaluable contribution in creating this resource, which was instrumental in making our work possible.

## Supplementary data

**FIG S1.**
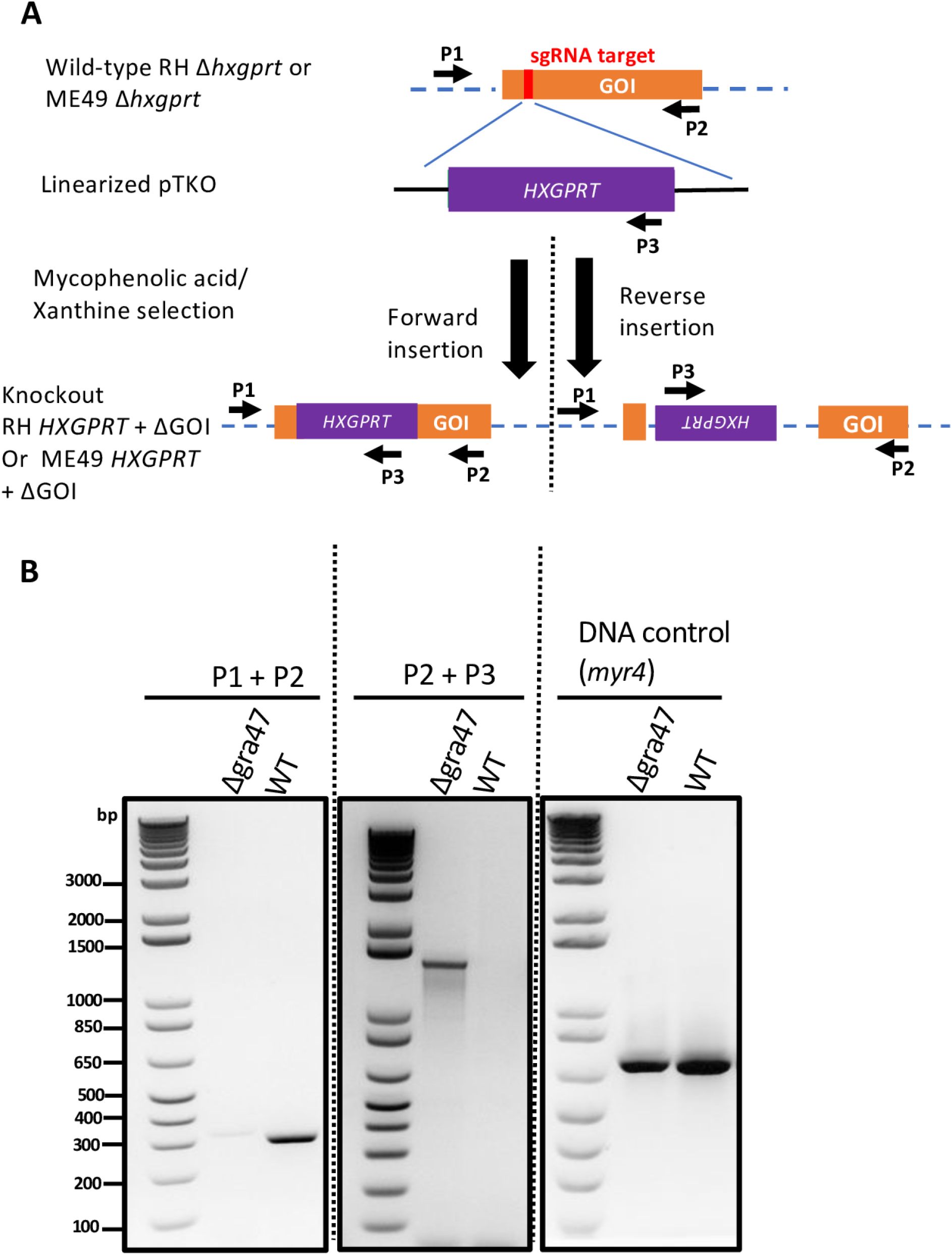
Generation of knockout parasite strains. **A)** The diagram illustrates the strategy employed to delete *GRA47* in both the type RH and ME49 strains. The CRISPR/Cas9-targeting site, depicted within a red box, indicates the specific region targeted for modification. To accomplish this, a linearized pTKO plasmid containing an *HXGPRT* selection cassette was utilized as a repair template. The selection process was conducted using mycophenolic acid and xanthine. **B)** Confirmation of the disruption of the gene of interest (GOI) was achieved using primers P1 and P2, which amplified a region within the GOI of 320 base pairs. For quality control of the PCR, amplification of MYR4 was used. To verify the successful insertion of the repair template, primers P2 and P3 were employed.

**FIG S2.**
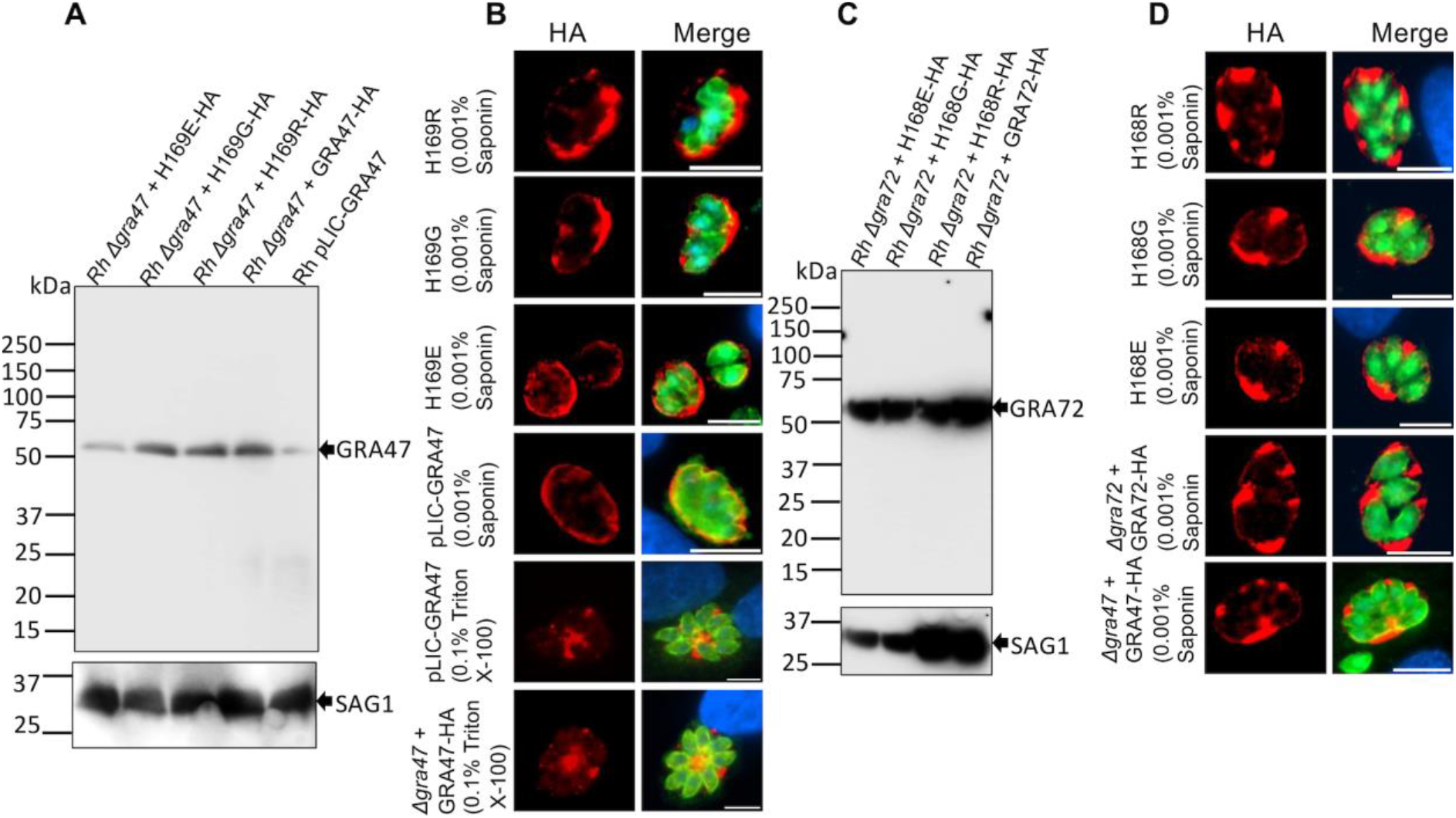
Complementation of Δ*gra47* and Δ*gra72* parasites with wild-type or histidine mutant derivatives. **A and C)** Shown is a Western blot demonstrating the successful complementation of the Δ*gra47* or Δ*gra72* knockout parasites in the type 1 (RH) background or C-terminally HA endotagged GRA47. SAG1 was used as the parasite loading control. Predicted molecular weights: GRA47= 51.54 kDa, SAG1= 34.83 kDa, GRA72= 57.51kDa. **B)** An immunofluorescence Assay (IFA) showing the localization of GRA47 from C-terminally HA-tagged or complemented Δ*gra47* parasites, marked in red, using different permeabilization methods. The scale bar represents 8μM. **D)** IFA showing the localization of Δ*gra72* parasites complemented with wild-type GRA72 or GRA72 containing histidine mutants. The scale bar used was 8μM.

**FIG S3.**
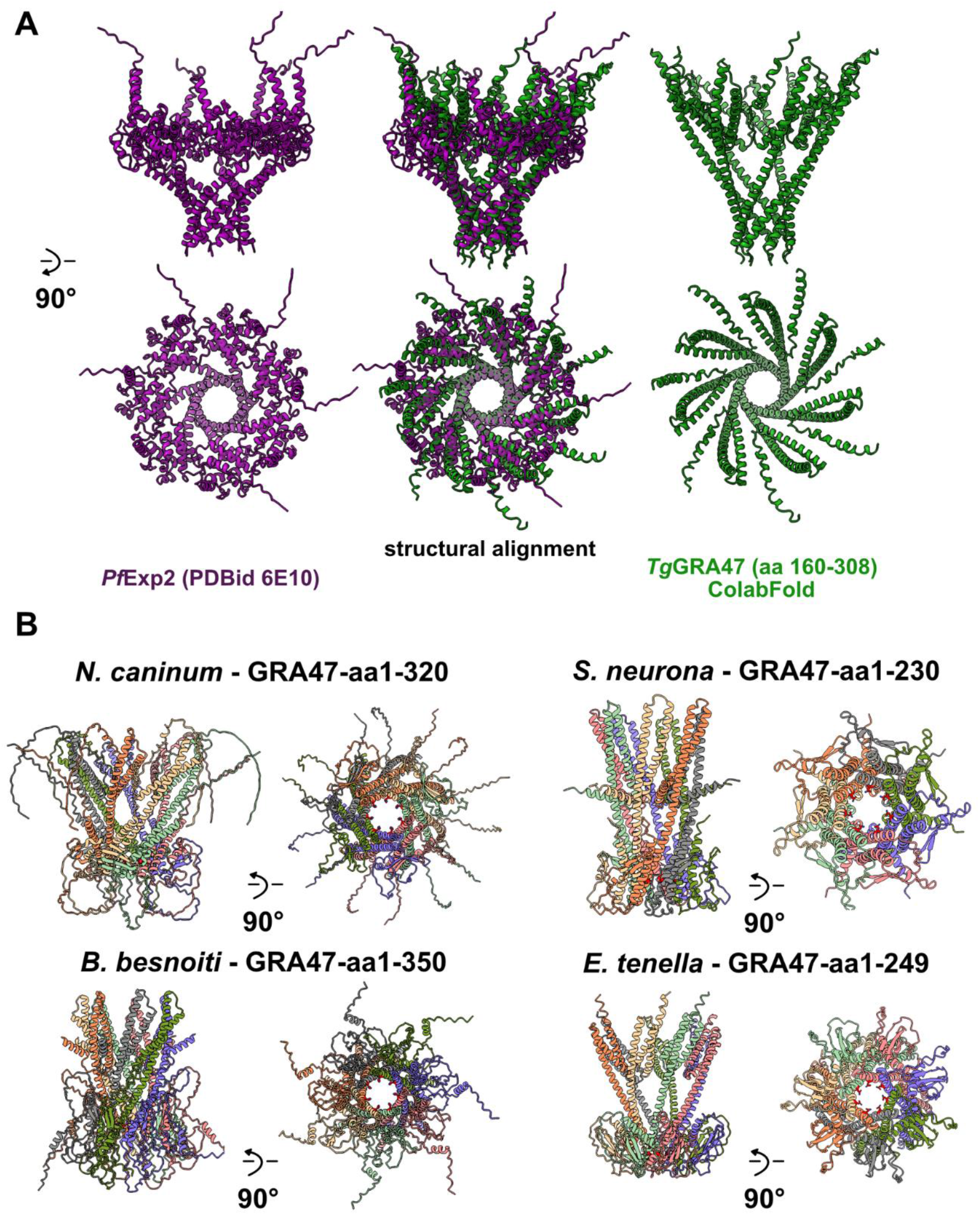
GRA47 orthologs are also predicted to form 7-mer pores. **A)** Comparison between the AlphaFold prediction of *Tg*GRA47 and the Exp2 7-mer pore cryo-EM structure. *Pf*Exp2 in magenta and the *Tg*GRA47 (aa 160 to 308) 7-mer model in green were structurally aligned on the N-terminal base of the transmembrane helix and displayed in a cartoon fashion. **B)** Pore forming orthologs of GRA47. Using the same approach as for *Tg*GRA47, *Neospora caninum*, *Sarcocystis neurona*, *Besnoitia besnoiti* and *Eimeria tenella* GRA47 orthologs determined by BLASTp search are also predicted to form 7-mer pore-like structures in AlphaFold-multimer predictions. Rank 1 models are displayed in a cartoon fashion with each monomer colored differently. The 90 ° rotation displays the pore entry with the gating residue at the constriction point shown in red.

**FIG S4.**
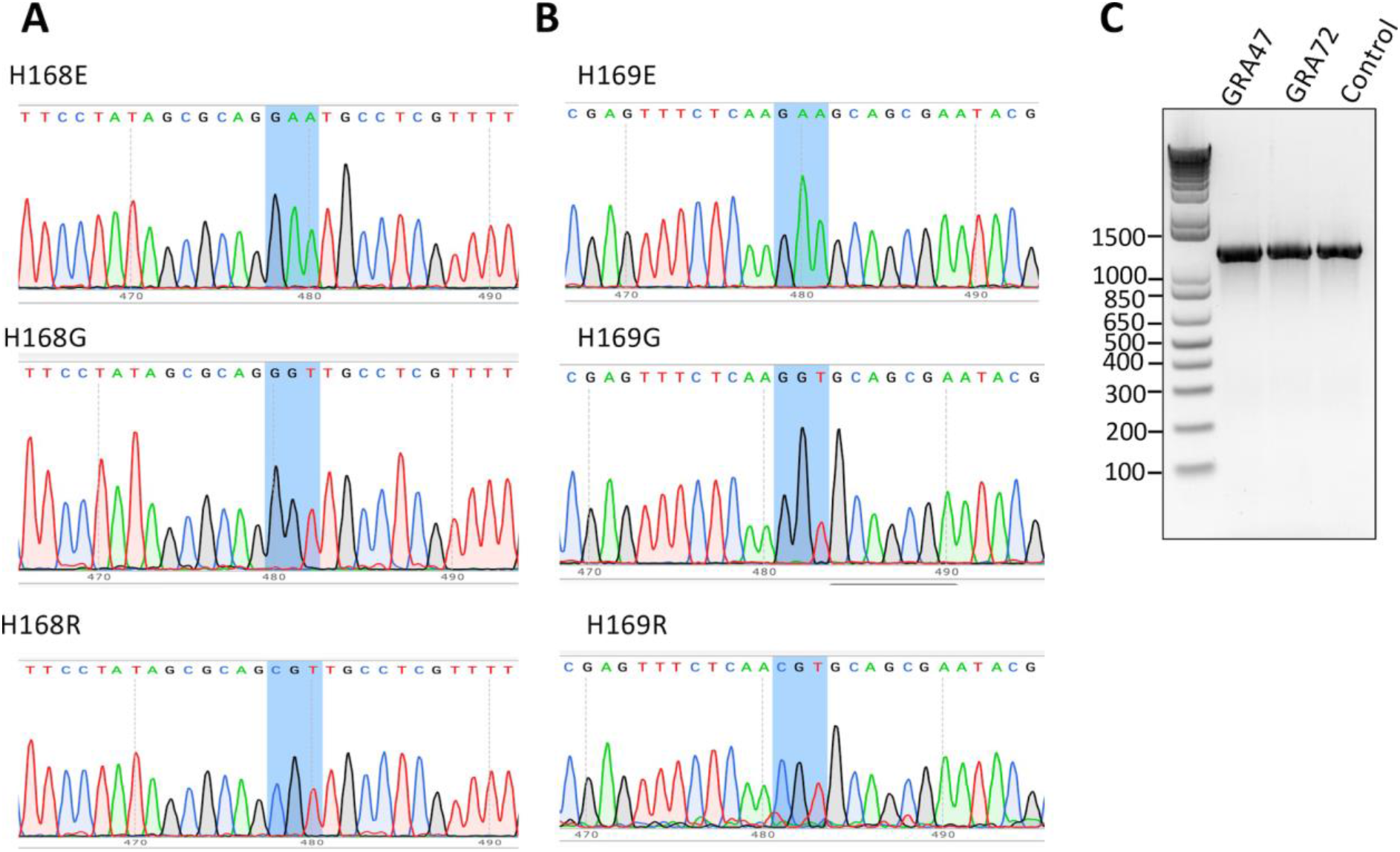
Confirmation of histidine mutations. Genomic DNA was extracted from **A)** GRA72 and **B)** GRA47 histidine mutant parasites and PCR amplification was performed. The PCR amplicon was Sanger sequenced. Shown are histidine mutations into glutamic acid (H168E and H169E), glycine (H168G and H169G) and arginine (H168R and H169R) of GRA72 and GRA47, respectively. **C)** Agarose gel electrophoresis showing the correct size of GRA47 and GRA72. Control is wild-type DNA without mutation.

**FIG S5.**
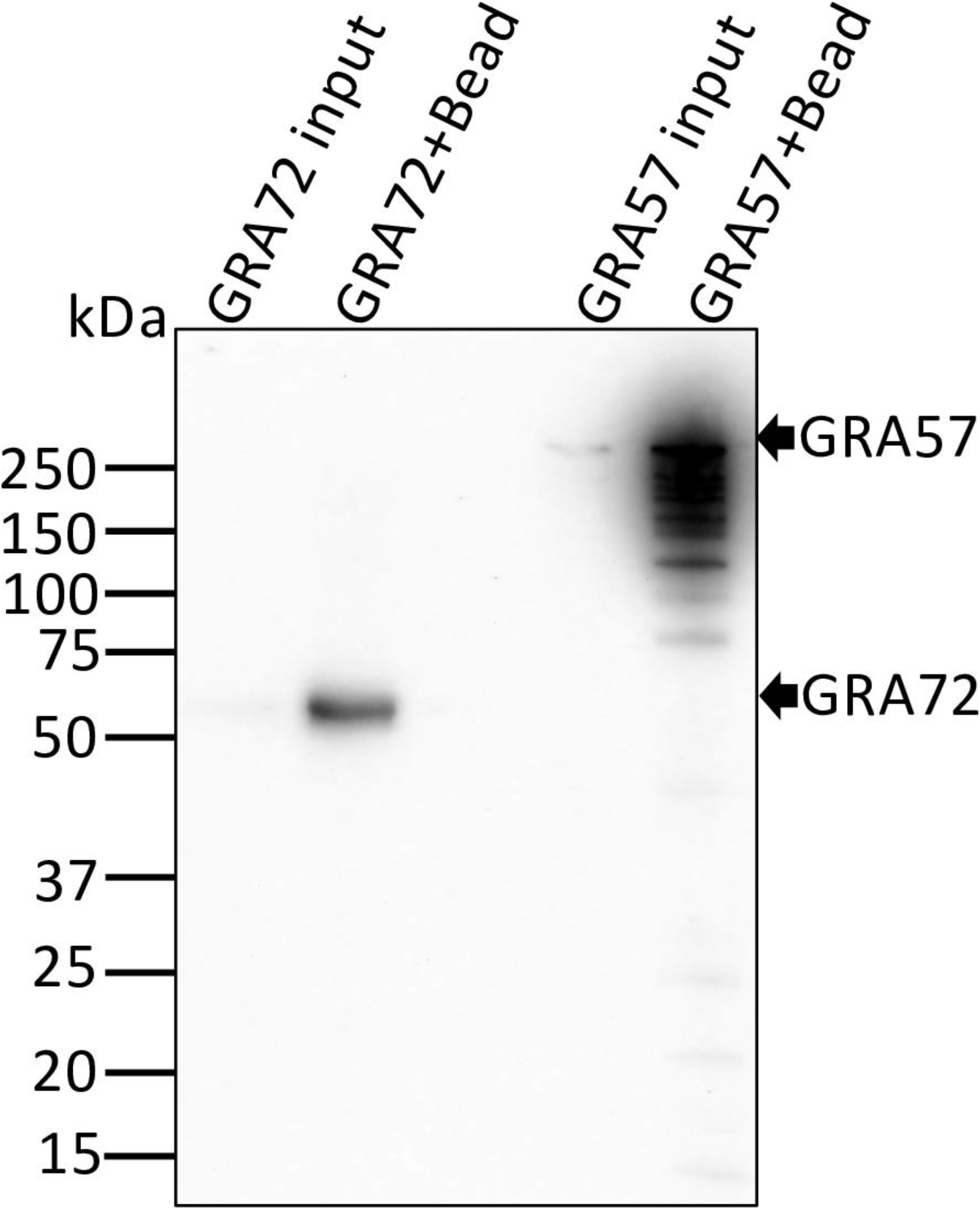
Pulldown of C-terminally HA-tagged GRA72 and GRA57. The Western blot, probed with an anti-HA antibody, displays the successful pulldown of C-terminally HA-tagged GRA72 and GRA57 from the type 1 (RH) background using anti-HA magnetic beads. Predicted molecular weight for GRA57 =246.19kDa and GRA72= 57.51kDa.

**FIG S6.**
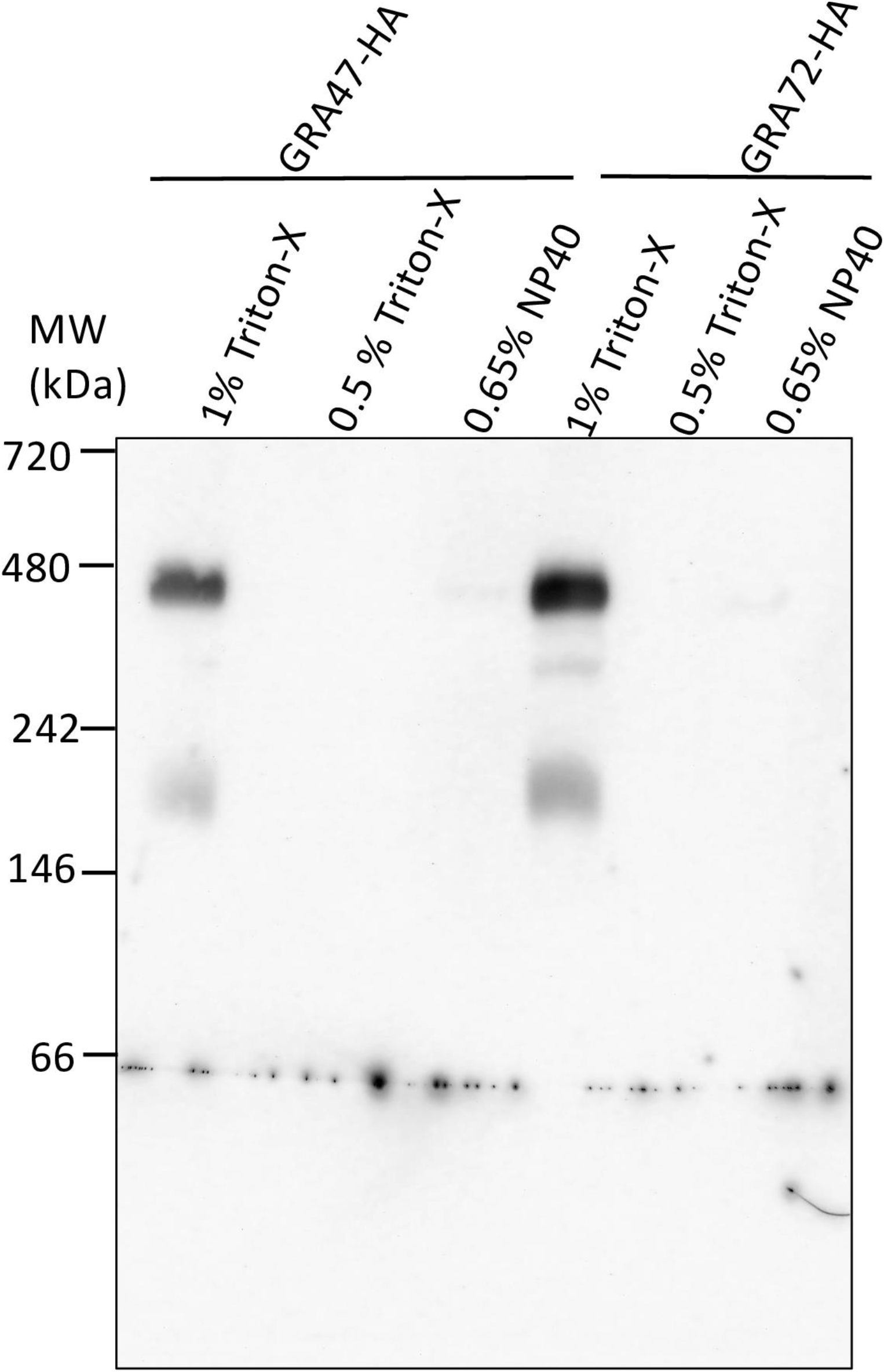
BluE-Native Page Electrophoresis of GRA47 and GRA72. A pellet collected from HFFs infected with the specified parasites prior to lysis underwent extraction with varying concentrations of detergents. Subsequently, the resultant samples were centrifuged, and the supernatants were loaded for blue native polyacrylamide gel electrophoresis (BN-PAGE). Following electrophoresis, Western blotting was conducted using an anti-HA antibody.

**FIG S7.**
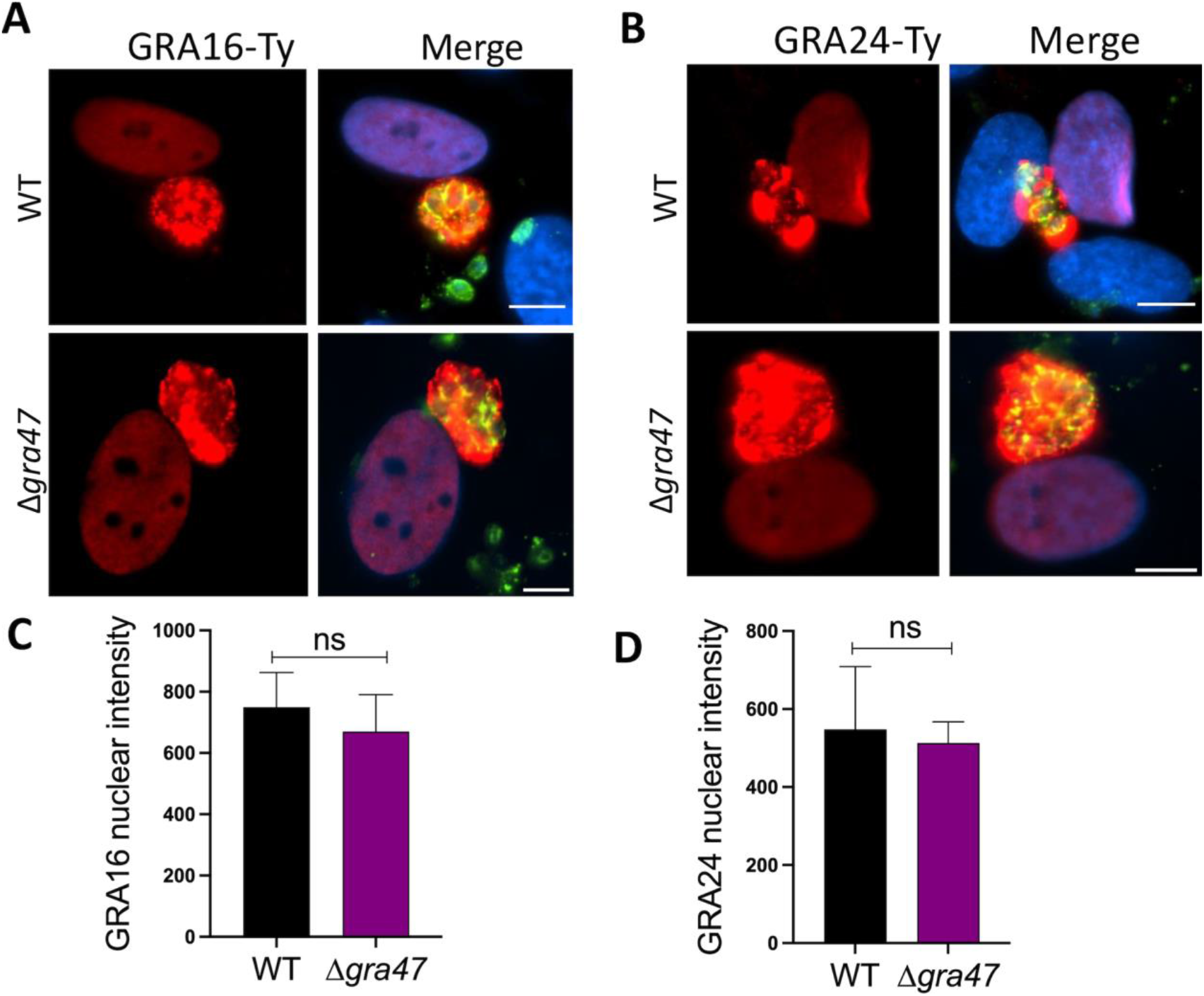
GRA47 does not affect export of GRA16 or GRA24 to the host cell. HFFs were infected with either wild-type (WT) or Δ*gra47* knockout parasites transiently expressing GRA16-Ty or GRA24-Ty. 24 hours post-infection, the cells were fixed with 3% formaldehyde and subsequently stained using a mouse anti-Ty antibody, shown in red. Panels **A** and **B** display representative images of GRA16 and GRA24 proteins being exported into the host nucleus, respectively. Panels **C** and **D** are the quantification of nuclear signal intensity for GRA16 and GRA24, respectively. Statistical analysis was conducted using a paired T-test (n=3). The images provided are representative of results obtained from three separate experiments, and the scale bars in the images indicate a length of 20 μm. ns= not significant.

